# CSF single-cell RNA sequencing reveals clonally expanded CD4^+^ stem cell-like memory T cells in GAD65-antibody associated neurological syndromes

**DOI:** 10.1101/2025.05.18.654720

**Authors:** Sumanta Barman, Saskia Räuber, Katharina Eisenhut, Daniela Esser, Martijn van Duijn, Madeleine Scharf, Marisol Herrera-Rivero, Paul Disse, Marius Jonas, Duygu Pul, Michael Heming, Louisa Müller-Miny, Manuela Paunovic, Christine Strippel, Ebru Haholu, Elisabeth Kaufmann, Justina Dargvainiene, Sabine Kahl, Marius Ringelstein, Eric Bindels, Heinz Wiendl, Nikolas H. Stoecklein, Johannes Fischer, Norbert Goebels, Lars Komorowski, Michael Roden, Andrea Rossi, Monika Stoll, Sven G. Meuth, Maarten J. Titulaer, Frank Leypoldt, Gerd Meyer zu Hörste, Franziska Thaler, Nico Melzer, cooperation with the EMC-AIE Study group, the German Network for Research on Autoimmune Encephalitis

**Affiliations:** Department of Neurology, Medical Faculty and University Hospital Düsseldorf, Heinrich Heine University Düsseldorf, Germany; Institute of Clinical Neuroimmunology, University Hospital, Ludwig-Maximilians-Universität München, Munich, Germany; Biomedical Center (BMC), Medical Faculty, Ludwig-Maximilians-Universität München, Martinsried, Germany; Graduate School of Systemic Neurosciences, Ludwig-Maximilians-Universität München, Munich, Germany; Institute of Clinical Chemistry, University Hospital Schleswig-Holstein Kiel/Lübeck, Germany; Department of Neurology, Erasmus MC University Medical Center, Rotterdam, the Netherlands; Institute for Experimental Immunology, affiliated to EUROIMMUN Medizinische Labordiagnostika AG, Lübeck, Germany; Institute of Epidemiology and Social Medicine, University of Münster, Münster, Germany; Department of Neurology with Institute of Translational Neurology, University Hospital Münster, Germany; Department of Neurology, LMU University Hospital, LMU Munich, Munich, Germany; Institute for Clinical Diabetology, German Diabetes Center (DDZ), Leibniz Institute for Diabetes Research at Heinrich-Heine-University, Düsseldorf, Germany; German Center for Diabetes Research (DZD), Partner Düsseldorf, München-Neuherberg, Germany; Department of Endocrinology and Diabetology, Medical Faculty and University Hospital Düsseldorf, Heinrich-Heine-University, Düsseldorf, Germany; Department of Neurology, Center for Neurology and Neuropsychiatry, LVR-Klinikum, Heinrich Heine University Düsseldorf, Düsseldorf, Germany; Department of Hematology, Erasmus MC University Medical Center, Rotterdam, the Netherlands; Department of General, Visceral and Pediatric Surgery, Medical Faculty and University Hospital Düsseldorf, Heinrich Heine University Düsseldorf, Germany; Institute for Transplantation Diagnostics and Cell Therapeutics, Medical Faculty and University Hospital Düsseldorf, Heinrich Heine University Düsseldorf, Germany; Genome Engineering and Model Development Lab (GEMD), IUF-Leibniz Research Institute for Environmental Medicine, Düsseldorf, Germany; Department of Genetic Epidemiology, Institute of Human Genetics, University of Münster, Münster, Germany; Department of Neurology, University Hospital Schleswig-Holstein, Kiel, Germany

## Abstract

**Background:** Glutamic acid decarboxylase (GAD) antibody-associated autoimmune neurological syndromes (AINS) are a spectrum of autoimmune-mediated CNS disorders. While antibodies targeting the 65 kDa isoform of GAD are of high diagnostic value, T cell mediated cytotoxicity has been identified as a key component of disease pathogenesis. The precise pathophysiological mechanisms by which the disease is triggered and maintained, however, remain incompletely understood.

**Methods:** We performed single-cell transcriptome and immune repertoire sequencing (sc-seq) in CSF and blood of 8 anti-GAD65 AINS patients compared to 8 non-inflammatory controls. Monoclonal antibodies (mAbs) were synthesized from B cell receptor (BCR) data to evaluate the B cellular immune response.

**Findings:** We identified an increase and expansion of activated CD4^+^ stem cell-like memory T cells (TSCM) in the CSF of anti-GAD65 AINS patients. Expanded T cells showed increased expression of proinflammatory genes. The mAb analysis revealed a high frequency of GAD65-reactive BCRs in the CSF of anti-GAD65 AINS patients with increased somatic hypermutations compared to non-GAD-reactive BCRs and BCRs from controls.

**Conclusions:** Sc-seq identified clonally expanded CD4^+^ TSCM in the CSF of anti-GAD65 AINS patients harboring cytotoxic properties likely contributing to disease pathogenesis. GAD-reactive B cells circulate in the CSF of anti-GAD65 AINS patients further supporting the concept of an antigen-specific intrathecal immune response. Future studies need to clarify the actual pathogenicity of these immune cells and the link between T and B cellular immune mechanisms in the pathogenesis of anti-GAD65 AINS.

**Funding:** German Research Foundation (ERARE18-202 UltraAIE), German Federal Ministry of Education and Research (CONNECT GENERATE (2.0); 01GM1908A and 01GM2208A).

## Introduction

Autoimmune neurological syndromes (AINS) with autoantibodies against the 65 kDa isoform of glutamic acid decarboxylase (GAD65) are immune-mediated disorders typically presenting with distinct clinical phenotypes: (1) limbic encephalitis (LE) with temporal lobe seizures (TLS) or epilepsy (TLE), (2) stiff-person-syndrome (SPS), (3) cerebellar ataxia (CA), or overlaps of the aforementioned ^1–3^. The disease course may be subacute or slowly progressing with limited treatment response often leading to lifelong neurological disability.

Autoantibodies are directed against GAD65, the rate-limiting enzyme of gamma-aminobutyric acid (GABA) synthesis. Given the cytoplasmic localization of the target antigen, the pathophysiological relevance of GAD65-antibodies remains controversial. While GAD65-antibodies are intrathecally produced in anti-GAD65 AINS patients and were shown to inhibit GAD65 enzymatic activity in vitro, neither a direct interaction between antibody and target antigen nor an antibody-mediated disease pathology could be demonstrated in animal models ^4,5^. Interestingly, recent neuropathological evidence from GAD-TLE patients highlights a pronounced infiltration of cytotoxic CD8^+^ T cells and plasma cells into the brain parenchyma ^6–8^. We recently reported a genome-wide association study (GWAS) of anti-GAD65 AINS, where the top genetic variant localized to a segment in the HLA class I region and several other variants with regulatory functions on gene expression mapped to CD4^+^ T cells and the cerebral cortex ^9^. These findings underline the complex immunopathogenesis of anti-GAD65 AINS and the need for systems biology approaches to unravel the concise pathophysiological mechanisms.

Here, we provide an in-depth characterization of peripheral and intrathecal immune cell populations in eight anti-GAD65 AINS patients compared to eight non-inflammatory controls, using single-cell RNA and immune repertoire sequencing. Our study identified a clonal expansion of CD4^+^ stem cell-like memory T cells (TSCM) with a cytotoxic phenotype in the CSF of anti-GAD65 AINS patients, which likely contribute to disease pathogenesis. Furthermore, we detected GAD-reactive B cells with an increased frequency of somatic hypermutations circulating in the CSF of these patients, suggesting they had undergone antigen-driven affinity maturation.

## Results

### Demographics and basic clinical data

Eight anti-GAD65 AINS patients were included in the study (**Figure 1A** and **Supplementary Table 1**). The median age of anti-GAD65 AINS patients was 65 [39 - 71] years. Seven patients were female, four presented with limbic encephalitis (LE), two with CA, one with SPS, and one showed an overlap of SPS and LE (**Figure 1B**). None of the patients received immunotherapy at the time of sampling. One patient had received steroids more than six months prior, and another had received immunoglobulins more than six years prior to sampling. Given the long interval between treatment and sampling, the patients can be considered immunotherapy-naïve ^10^. Three patients showed CSF pleocytosis at sampling. Blood-CSF-barrier dysfunction (BCSFBD) was only apparent in one patient. Three patients had CSF-specific oligoclonal bands (OCBs) (**Supplementary Table 2**).

**Figure 1.**
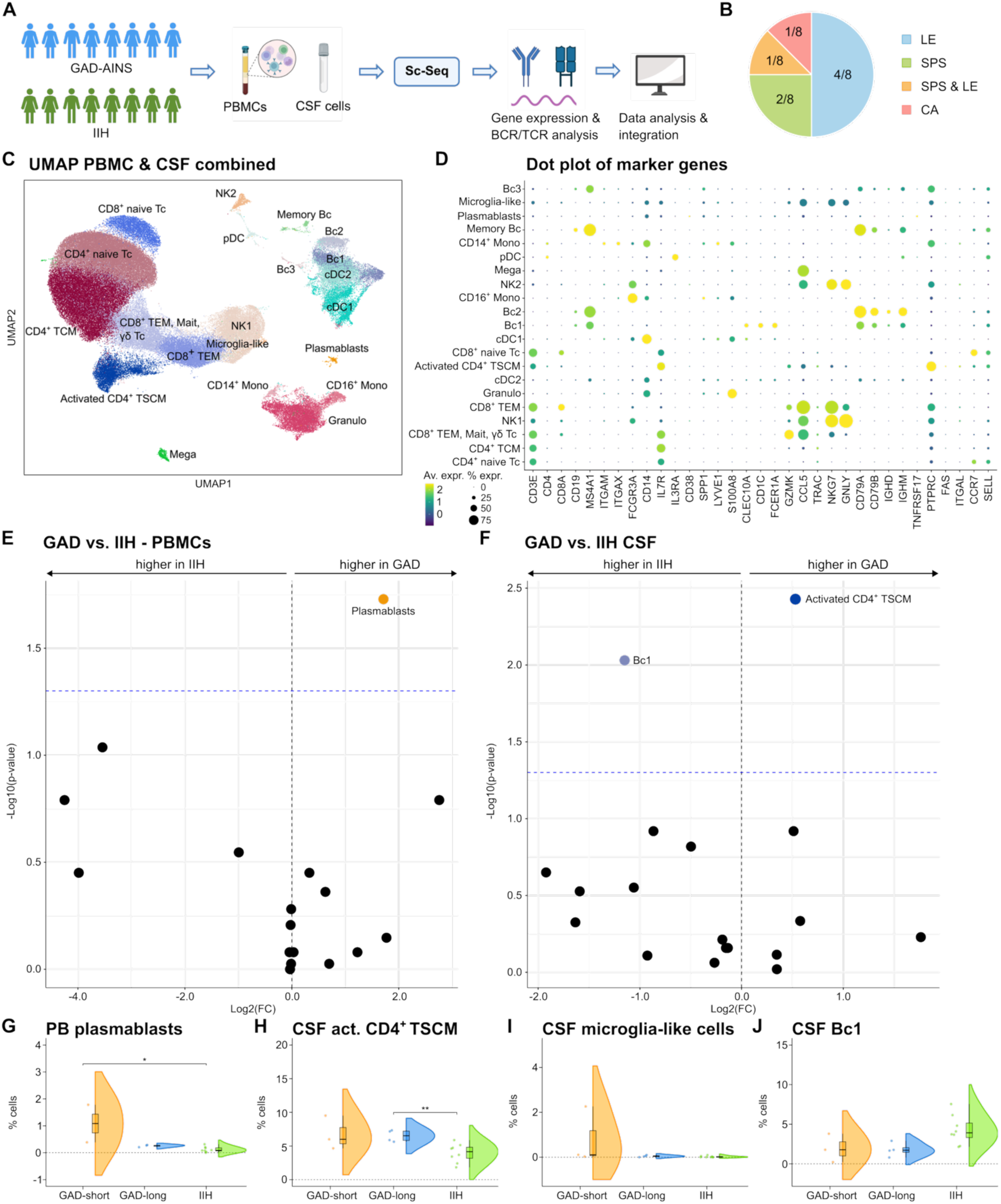
CD4^+^ TSCM and microglia-like cells are expanded in the CSF and plasmablasts in the PB of anti-GAD65 AINS patients. **A** Study design. Created in BioRender. Räuber, S. (2025) https://BioRender.com/4ir3vf0; **B** Clinical phenotypes of anti-GAD65 AINS patients. Number of patients with each clinical phenotype in relation to the total number of patients are shown. **C** UMAP showing 21 color-coded cell clusters of 1,18,492 single cell transcriptomes integrated from CSF cells and PBMC from anti-GAD65 AINS patients (n = 8) and IIH patients (n = 8). **D** Dot plot illustrating selected marker genes of cell clusters from CSF cells and PBMC from anti-GAD65 AINS and IIH patients. Color encodes mean expression, dot size visualizes the fraction of cells expressing the gene. **E, F** Volcano plots depicting differentially abundant PB (**E**) and CSF (**F**) cell clusters in anti-GAD65 AINS compared to IIH patients. The log2 fold change of the differential abundance is plotted against the negative log10 of the p-values assessed by Wilcoxon signed-rank test. The threshold for the p-value was set at 0.05. The microglia-like cell cluster is not plotted given the median of 0 in the IIH group. **G-J** Violin plots with overlaying box plots depicting the differences in PB plasmablasts (**G**), CSF activated CD4^+^ TSCM (**H**), CSF microglia-like cells (**I**), and CSF Bc1 (**J**) between anti-GAD65 AINS with short disease duration (< 1 year), anti-GAD65 AINS with long disease duration (> 1 year), and IIH patients. Boxes display the median as well as the 25th and 75th percentiles. The whiskers extend from the hinge to the largest and smallest values, respectively, but no further than 1.5 * IQR from the hinge. P-values were calculated by Kruskal Wallis test with Dunn post hoc test (p-adjustment method: Benjamini–Hochberg). * p ≤ 0.05, ** p ≤ 0.01. *Act. - activated; Av - average; Bc - B cells; BCR - B cell receptor; CA - cerebellar ataxia; CSF - cerebrospinal fluid; cDC - conventional dendritic cell; DC - dendritic cells; expr. - expression; FC - fold change; GAD - patients with glutamic acid decarboxylase antibody-associated autoimmune neurological syndromes (AINS); Granulo - granulocytes; IIH - idiopathic intracranial hypertension; LE - limbic encephalitis; long - long disease duration (> 1 year); MAIT - Mucosal-associated invariant T cells; Mega - megakaryocytes; Mono - monocytes; NK - natural killer cells; PB - peripheral blood; PBMC - Peripheral Blood Mononuclear Cell; pDC - plasmacytoid dendritic cells; short - short disease duration (< 1 year); Tc - T cells; TCR - T cell receptor; Sc-Seq - single-cell sequencing; SPS - stiff-person syndrome; TCM - central memory T cells; TEM - effector memory T cells; TSCM - stem cell-like memory T cells; UMAP - Uniform Manifold Approximation and Projection*.

In addition, eight patients with Idiopathic Intracranial Hypertension (IIH) were included as non-inflammatory controls (**Figure 1A** and **Supplementary Table 3**). The median age of the IIH patients was 38 [27-52] years and seven were female. Two patients had CSF pleocytosis and/or BCSFBD. None of these patients received immunotherapy at the time of or prior to sampling. Basic CSF characteristics of the IIH cohort are displayed in **Supplementary Table 4**.

### Increase in peripheral blood (PB) plasmablasts and CSF activated CD4^+^ TSCM in anti-GAD65 AINS patients

Applying single-cell RNA sequencing (scRNA-seq), we analyzed peripheral blood mononuclear cells (PBMCs) and CSF cells in an unbiased manner. We integrated scRNA-seq data from PBMC and CSF of anti-GAD65 AINS and IIH patients, yielding an object encompassing 118,492 total single-cell transcriptomes (PBMCs: 63,176, CSF: 55,316) with 4231.8 ± 691.5 (mean ± SEM) cells per sample and 1280.1 ± 1.65 (mean ± SEM) genes detected per cell. Hereafter, we refer to single-cell transcriptomes as ‘cells’. Then, we performed unsupervised clustering, which resulted in 21 cell clusters (**Figure 1C**). Cluster annotation was based on marker gene expression using a combination of manual and automated approaches (**Figure 1D**). When comparing anti-GAD-AINS to IIH patients, we found an increased proportion of plasmablasts in the PB of anti-GAD65 AINS patients (**Figure 1E**). In the CSF, the B cell (Bc) cluster 1 was reduced, while there was an expansion of the activated CD4^+^ TSCM cluster in anti-GAD65 AINS identified by the expression of typical TSCM markers (e.g., PTPRC (CD45-RA), FAS, ITGAL, CCR7, SELL, IL7R) ^11,12^ (**Figure 1F**). Furthermore, higher percentages of microglia-like cells (MLC) were found in the CSF of anti-GAD65 AINS compared to IIH patients. We next subdivided the anti-GAD65 AINS cohort in patients with short disease duration (< 1 year; n = 3) and long disease duration (> 1 year; n = 5) at the time of sampling and compared the significant cell clusters between those two subgroups and IIH controls. PB plasmablasts were especially increased in anti-GAD65 AINS patients with short disease duration (**Figure 1G**) while CSF activated CD4^+^ TSCM were significantly higher in anti-GAD65 AINS patients with long disease duration in comparison to IIH patients (**Figure 1H**). No relevant differences were noted with regard to CSF MLC and CSF Bc1 between groups (**Figure 1I, J**).

In summary, we detected a higher fraction of PB plasmablasts in anti-GAD65 AINS compared to IIH patients, especially in individuals with short disease duration, while activated CD4^+^ TSCM were increased in the CSF of anti-GAD65 AINS in comparison to IIH patients. This increase was more prominent in anti-GAD65 AINS patients with long disease duration.

### Differences in gene expression involved in the regulation of inflammatory immune responses between anti-GAD65 AINS and IIH patients

In order to identify changes in gene expression patterns in anti-GAD65 AINS patients in comparison to IIH patients, we performed differential gene expression (DGE) analysis and created volcano plots of differentially expressed genes (**Figure 2A-E**). First, CSF and PBMC analysis was performed across all cell types, respectively (**Figure 2A, B**). Next, the analysis was repeated for all PBMC and CSF cell clusters separately (**Figure 2C-F**).

**Figure 2.**
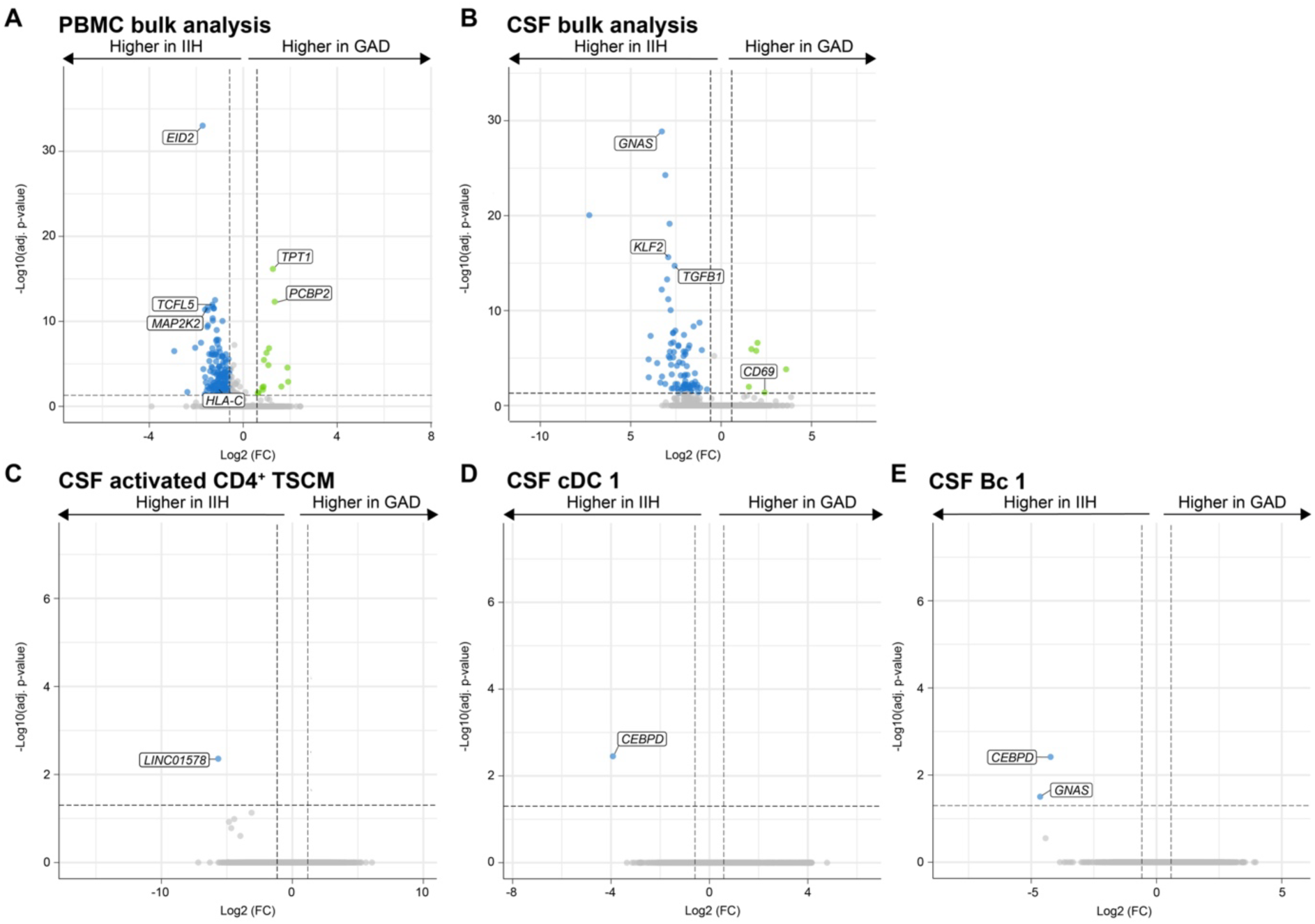
Differentially expressed genes involved in regulation of the immune response between anti-GAD65 AINS and IIH patients. **A-E** Volcano plots depicting the differentially expressed genes in the PB and CSF of anti-GAD65 AINS compared to IIH patients. Pseudobulking was performed and the differentially expressed genes were calculated using the ‘FindMarkers’ function of Seurat. Adjusted p<0.05 and log2FC≥0.585 were used as cut-offs: **A, B** PBMC and CSF bulk analysis, respectively. **C** DGE analysis of CSF activated CD4^+^ TSCM. **D:** DGE analysis of CSF cDC cluster 1. **E** DGE analysis of CSF B cell cluster 1. *Adj. - adjusted; Bc - B cells; CEBPD - CCAAT/enhancer-binding protein delta; CSF - cerebrospinal fluid; DC - dendritic cells; EID2 - EP300-interacting inhibitor of differentiation 2; FC - fold change; GAD - patients with glutamic acid decarboxylase antibody-associated autoimmune neurological syndromes (AINS); GNAS - Neuroendocrine secretory protein 55; HLA-C - Human Leukocyte Antigen-C; IIH - idiopathic intracranial hypertension; KLF2 - Krueppel-like factor 2; LINC01578 - Long noncoding RNA LINC01578; MAP2K2 – Dual specificity mitogen-activated protein kinase kinase 2; PB - peripheral blood; PBMC – Peripheral Blood Mononuclear Cell; PCBP2 - Poly(rC)-binding protein 2; TCFL5 - Transcription factor-like 5 protein; TGFB1 - transforming growth factor-β; TPT1 - Translationally-controlled tumor protein; TSCM - stem cell-like memory T cells*.

Among the top differentially expressed genes in the PBMC bulk analysis were *EID2 -* which functions as a chemo-attractant receptor mediating cell migration ^13^ -, *TCFL5* - which was previously linked to impaired germinal center formation, B cell differentiation, and B cell receptor (BCR) signaling ^14^ -, *MAP2K2* - reported to modulate expression of MHCII, CD40, CD80 and to have anti-inflammatory effects ^15,16^ -, *TPT1* - which was found to be increased in twins with multiple sclerosis (MS) compared with unaffected twins ^17^ -, and *PCBP2* - essential for CD4^+^ T cell activation and proliferation ^18^. Furthermore, one gene, which we found to be associated with anti-GAD65-AINS in a genome-wide association study, was downregulated in the current anti-GAD65 AINS cohort (*HLA-C*) ^9^ (**Figure 2A**). Regarding the CSF, the following genes were among the top differentially regulated ones: *GNAS* - a gene which was previously associated with tumor immune cell infiltration and Th2 polarization ^19,20^ -, *KLF2* - a transcription factor critical for immune cell differentiation, activation, proliferation, and migration as well as generation of Tregs ^21–23^ -, *CD69* - a classical marker of lymphocyte activation and tissue retention and regulator of Treg differentiation ^24^ -, and *TGFB1* - a cytokine critical for both immune tolerance and immunity ^25^ (**Figure 2B**). Having a closer look at the different cell clusters, we found the expression of *LINC01578* - encoding a long non-coding RNA (lncRNA) ^26^ - to be lower in activated CD4^+^ TSCM of anti-GAD65 AINS in comparison to IIH patients (**Figure 2C**). Conventional dendritic cells (cDC) 1 and Bc1 of anti-GAD65 AINS patients had reduced *CEBPD* expression - known to be implicated in regulation of IL-10 expression ^27^ - compared to IIH patients. Bc1 of anti-GAD65 AINS patients also showed decreased *GNAS* expression (**Figure 2 D-E**).

In summary, DGE analysis identified differences in the expression of several genes involved in the regulation of the inflammatory immune response between anti-GAD65 AINS and IIH patients. These might present therapeutic targets warranting further investigation.

### Numerous CSF BCRs are GAD65-reactive and somatically hypermutated, but originate from non-expanded B cells

We next sought to characterize the intrathecal B cell response in anti-GAD65 AINS patients. BCR sequence analysis identified little to no clonally expanded B cells in the CSF compartment of anti-GAD65 AINS patients (**Supplementary Figure 1A-C**). To determine whether those BCRs were still autoantigen-specific, we randomly selected CSF BCRs of different Ig isotypes with concurrently available consensus heavy and light chain BCR variable region sequence information from a representative subgroup of patients (n=4, **Figure 3A**). The corresponding heavy and light chain sequences were synthesized, cloned, expressed, and affinity purified as recombinant monoclonal antibodies (mAbs) (**Figure 3A)**. Remarkably, out of 40 mAbs, 10 (25%) were GAD65-reactive, as shown by indirect immunofluorescence staining of transfected HEK293T cells (**Figure 3B-D, Supplementary Figure 1D**) and non-human primate brain tissue (**Supplementary Figure 1E**). Notably, 3 out of 10 GAD65-reactive mAbs additionally showed Purkinje-cell reactivity on non-human primate brain tissue stainings (**Supplementary Figure 1E**). None of the 40 mAbs displayed any other clear anti-CNS reactivity. GAD65-reactive BCRs that could be assigned to specific B cell subsets (3/10) were mapped to memory and naïve B cell clusters. While immunoglobulin isotype distribution did not differ between anti-GAD65 AINS and IIH patients in the CSF (**Supplementary Figure 1F**), GAD65-reactive mAbs were exclusively of the IgG1 isotype and significantly more somatically hypermutated than non-GAD65-reactive mAbs (**Figure 3C, E**), suggesting that they likely have undergone antigen-driven affinity maturation. In general, significantly more somatic hypermutations were found in the CSF BCRs of anti-GAD65 AINS compared to IIH patients (**Figure 3F**).

**Figure 3.**
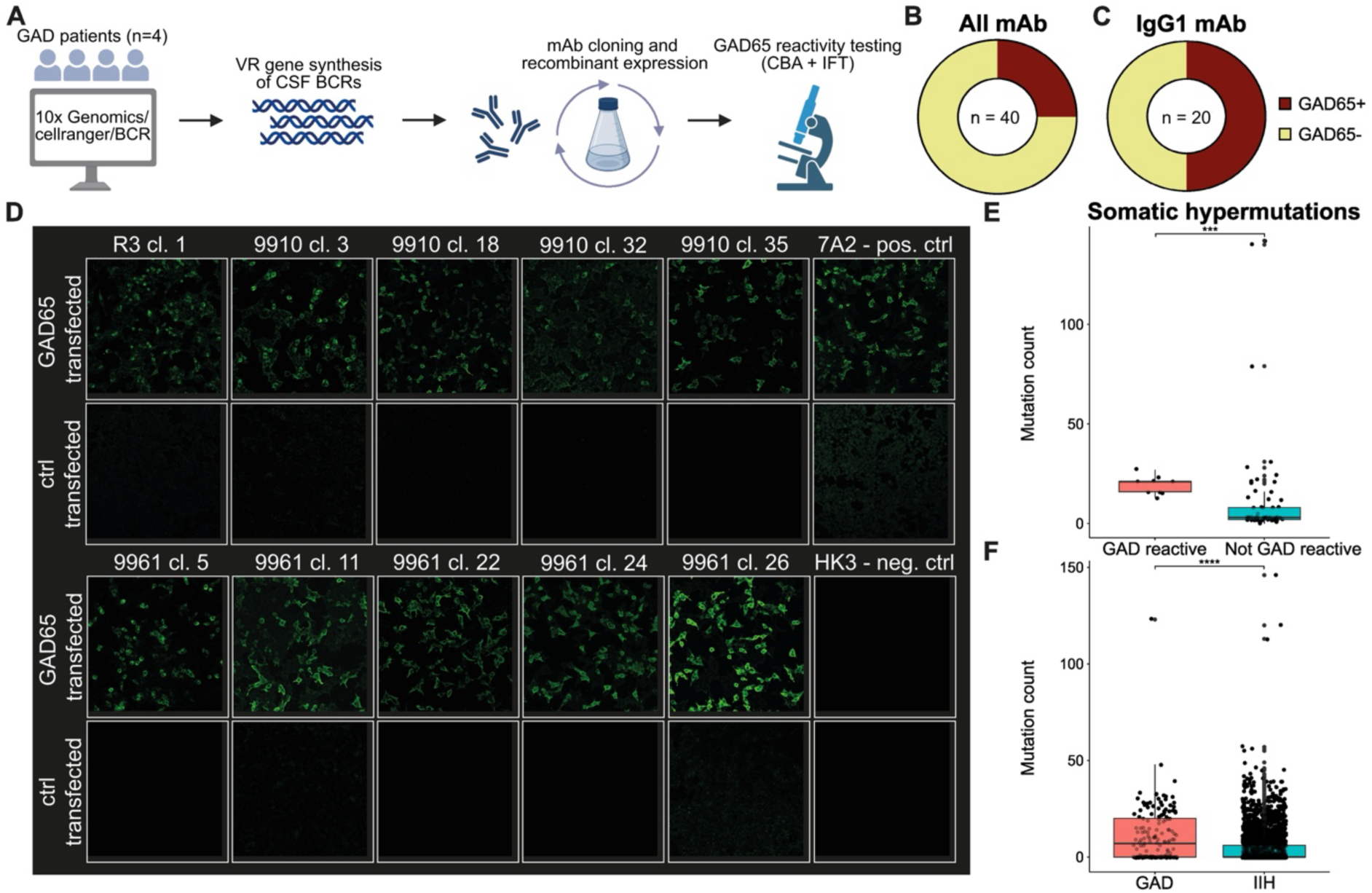
Recombinant production of monoclonal antibodies from CSF BCR data reveals numerous GAD65-reactive BCRs with increased somatic hypermutations. **A** Monoclonal antibody production workflow: 40 recombinant mAbs were produced from n=4 GAD patients by gene synthesis of CSF BCR variable region sequences of matching heavy and light chains as extracted from Cell Ranger-processed data, subsequent cloning, expression, and affinity purification. GAD65-reactivity testing by cell-based assay and indirect immunofluorescence stainings on primate brain tissue showed GAD65-reactivity in **B** 10 out of 40 produced mAbs (25%) and **C** 10 out of 20 produced IgG1 mAbs (50%). **D** Immunofluorescence stainings of the 10 GAD65-reactive recombinant human mAbs (green) to HEK cells overexpressing GAD65 or control-transfected cells (as indicated in row caption). Each column represents a tested mAb (as indicated by patient ID and clonotype number in column caption). 7A2 serves as positive and HK3 as negative control, both monoclonal antibodies which were produced in the same expression vector. **E** Quantification of somatic hypermutations in produced GAD65-reactive and non-reactive mAbs and **F** all CSF BCRs from anti-GAD65 AINS as compared to IIH patients. *BCR - B cell receptor; CBA - cell-based assay; cl. - clonotype; GAD - patients with glutamic acid decarboxylase antibody-associated autoimmune neurological syndromes (AINS); GAD65+ - glutamic acid decarboxylase 65-reactive; GAD65- - not glutamic acid decarboxylase 65-reactive; IFT - indirect immunofluorescence testing; IIH - idiopathic intracranial hypertension; CSF - cerebrospinal fluid; mAb - monoclonal antibody; neg. ctrl - negative control; pos. ctrl - positive control; VR - variable region*.

In summary, anti-GAD65 AINS patients harbor numerous GAD65-reactive and somatically hypermutated BCRs in the CSF compartment, which however originate from non-expanded B cells.

### Compartment-specific TCR clonal expansion in anti-GAD65 AINS

To analyze the T cell receptor (TCR) repertoire, we used IgBlast-aligned V(D)J data from T cells, combined with transcriptomic data from T cell clusters. Analysis of TCR repertoires revealed distinct patterns of clonal expansion between anti-GAD65 AINS and IIH patients across different compartments. In the CSF of anti-GAD65 AINS patients, we observed significantly expanded T cell clones characterized by unique TCR sequences, suggesting an antigen-driven immune response (**Figure 4A-F**). These expanded CSF clones demonstrated a distinct transcriptional profile, with DGE analysis revealing significant upregulation of genes involved in cellular cytotoxicity, TCR signaling, T cell proliferation and survival, chemotaxis, as well as in inflammatory immune responses (**Figure 4B-C**). In contrast, the CSF compartment of the control group showed limited clonal expansion, indicating a more diverse and non-targeted immune repertoire (**Figure 4D-E**). Intriguingly, this pattern was reversed in the PBMC compartment, where control subjects exhibited higher frequencies of expanded clones compared to anti-GAD65 AINS patients (**Figure 4D-E**). Analysis of the top 20 most abundant TCR clones in the CSF further confirmed this pattern, demonstrating elevated clonal frequencies in anti-GAD65 AINS compared to IIH patients suggesting a focused, disease-specific immune response (**Supplementary Figure 2A-B**). Again, this pattern was inverted in the PBMC compartment, where control subjects displayed higher frequencies of expanded clones compared to anti-GAD65 AINS patients (**Supplementary Figure 2A-B**). Furthermore, TCR clonal overlap was observed between PBMC and CSF compartments in both anti-GAD65 AINS and IIH groups. Notably, a higher degree of clonal sharing was evident in the majority of IIH samples (5 out of 6) compared to anti-GAD65 AINS samples (2 out of 4). Moreover, a limited number of TCR clones were found to be shared across different samples in both groups (**Supplementary Figure 2 C-D)**.

**Figure 4.**
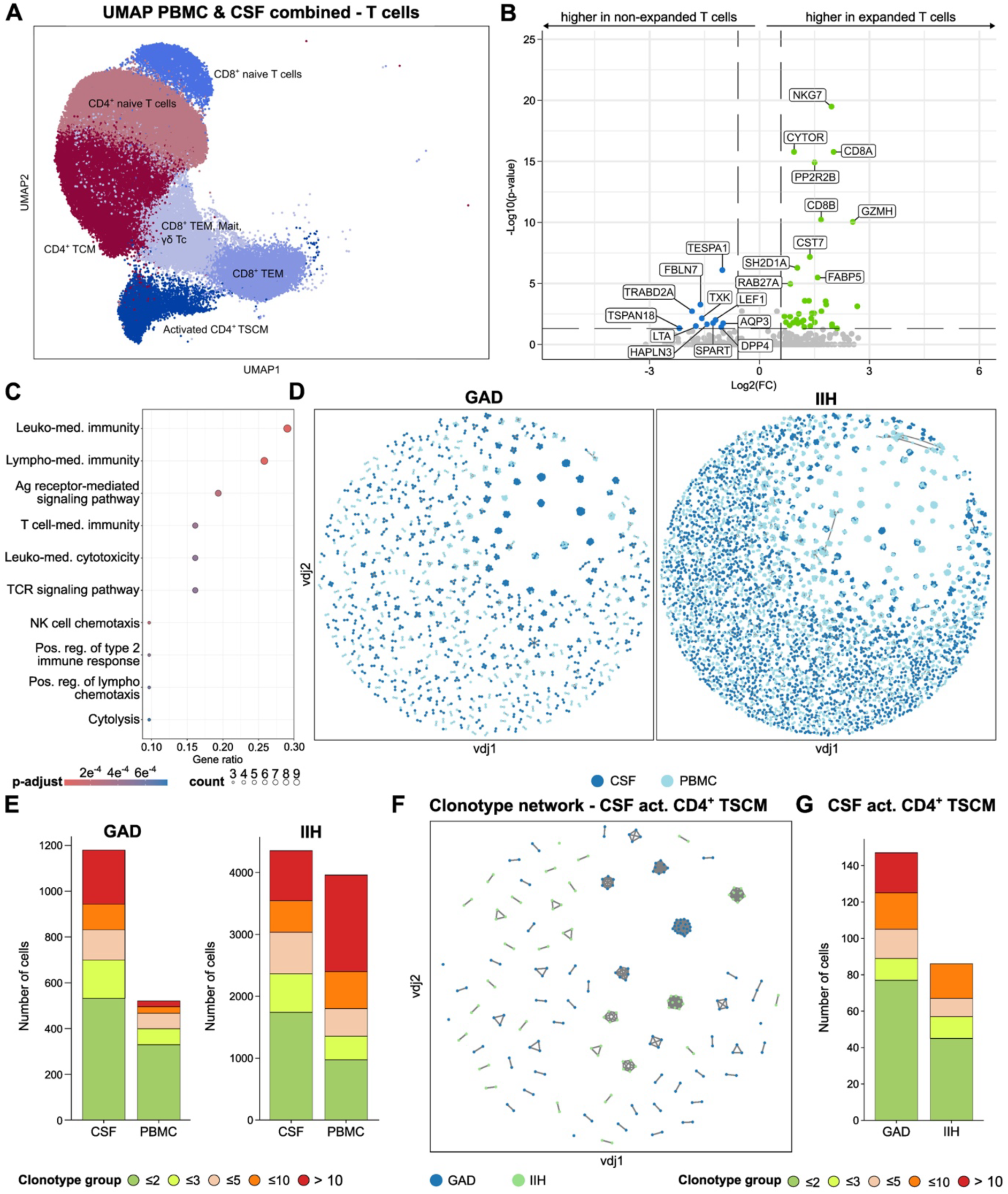
Comprehensive analysis of T cell repertoire in anti-GAD65 AINS patients and controls reveals distinct clonal expansion patterns. **A** UMAP visualization of T cell repertoire composition of anti-GAD65 AINS and IIH patients. **B** Pseudobulk differential gene expression analysis displayed as a volcano plot comparing expanded versus non-expanded T cell clones in CSF of anti-GAD65 AINS patients. Adjusted p<0.05 and log2FC≥ 0.585 were used as cut-offs. Top 10 up-regulated and down-regulated genes are labelled. **C** Gene Ontology (GO) enrichment analysis of differentially expressed genes in expanded CSF T cell clones from anti-GAD65 AINS patients. Dot size represents gene count; color intensity indicates statistical significance (adjusted p-value). **D** Clonal network visualization depicting TCR repertoire relationships between CSF and PBMC compartments in anti-GAD65 AINS and IIH patients. Node size indicates clonal frequency; edges represent shared TCR sequences. **E** Stacked bar plot represents quantitative comparison of TCR clonal distribution in expanded clonal groups, comparing CSF and PBMC compartments in anti-GAD65 AINS and IIH patients. Each stack shows the relative frequency of a distinct expanded clonal group. **F** Comparative clonal network analysis of CD4^+^ TSCM cells in CSF compartments between anti-GAD65 AINS and IIH patients, highlighting disease-specific repertoire patterns. Node size indicates clonal frequency; edges represent shared TCR sequences. **G** Stacked bar plot represents quantitative comparison of CD4^+^ TSCM cell TCR clonal compositions in expanded clonal groups from CSF, contrasting anti-GAD65 AINS with IIH patients. Each stack represents the relative frequency of a distinct expanded clonal group. *Act. - activated; Ag - antigen; UMAP - Uniform Manifold Approximation and Projection; CSF - cerebrospinal fluid; GAD - patients with glutamic acid decarboxylase antibody-associated autoimmune neurological syndromes (AINS); IIH - idiopathic intracranial hypertension; FC - fold change; GO - Gene Ontology; Leuko - leukocyte; med. - mediated; NK - natural killer cells; pos. - positive; reg. - regulation; TCM - central memory T cells; TCR - T cell receptor; TEM - effector memory T cells; TSCM - stem cell-like memory T cells; PBMC - Peripheral Blood Mononuclear Cell*.

In addition to the T cell clonal expansion analysis, which identified expansion of specific T cell clones in the CSF of anti-GAD65 AINS patients, indicating proliferation in response to disease-relevant antigens, we went on and performed TCR diversity analysis to assess the overall composition and variability of the TCR repertoire. We found a significantly reduced diversity in the PBMC compartment of anti-GAD65 AINS compared to IIH patients, consistent with a migration of pathogenic T cell clones to the CSF (**Supplementary Figure 2E**). However, no significant differences in TCR diversity were observed in the CSF compartment between the two groups (**Supplementary Figure 2F**).

### Expansion of CD4^+^ TSCM cells in the CSF of anti-GAD65 AINS patients

Detailed characterization of CD4^+^ TSCM cells in the CSF revealed distinct clonal expansion patterns between anti-GAD65 AINS and IIH patients. Analysis of the TCR repertoire demonstrated a marked enrichment of expanded clones within the CSF CD4^+^ TSCM cell population of anti-GAD65 AINS compared to IIH patients (**Figure 4F, G**). This pattern was particularly evident in the clonal network analysis, where anti-GAD65 AINS patients exhibited larger, interconnected clusters of clonally expanded TSCM cells (frequency ≥2), suggesting the presence of antigen-driven clonal selection (**Figure 4F**). Quantitative assessment of clonal distributions further supported this observation. Anti-GAD65 AINS patients showed a higher frequency of expanded clonal groups within their CSF TSCM compartment (**Figure 4G**). The disease-specific nature of this expansion was further emphasized by individual clonotype network analysis, which revealed distinct architectural patterns in anti-GAD65 AINS patients characterized by larger nodes representing highly expanded clones (**Supplementary Figure 3A-B**). Analysis of the top 20 most abundant TCR clones in the CSF TSCM population demonstrated higher clonal frequencies in anti-GAD65 AINS compared to IIH patients, indicating a more focused and potentially disease-relevant immune response within this memory T cell subset (**Supplementary Figure 3C**).

## Discussion

Our study characterizes the immune landscape of the CSF and PB of anti-GAD65 AINS patients at single-cell resolution. We report several important aspects: (1) The fraction of CD4^+^ TSCM cells was increased in the CSF of anti-GAD65 AINS patients, especially in individuals with long disease duration, while plasmablasts were increased in the PB, particularly in those with short disease duration. (2) CSF CD4^+^ TSCM cells were clonally expanded in anti-GAD65 AINS patients, while peripheral TCR diversity was reduced. (3) Clonally expanded T cells in the CSF of anti-GAD65 AINS patients showed a proinflammatory gene signature. (4) CSF BCR repertoire analysis revealed a relevant proportion of GAD65-reactive BCRs that belong to the IgG1 subtype and were derived from non-clonally expanded cells.

We suggest that the markedly increased frequencies of activated and clonally expanded CD4^+^ TSCM in CSF of anti-GAD65 AINS patients are disease relevant. Memory T cells are an important part of the adaptive immune system, providing long-term protection against pathogens. When they re-encounter an antigen, they proliferate rapidly and mediate cellular cytotoxicity ^28^. Memory T cells have been implicated in several autoimmune disorders, including MS and type 1 diabetes (T1D). In MS, an increase in activated effector memory CD8^+^ T cells and oligoclonal expansion of memory CD8^+^ T cells has been detected in the CSF ^29,30^. Within the brain parenchyma, an invasion of tissue-resident memory T cells was noted ^31^. Myelin-reactive T cells in MS patients were found more often in the memory population, while those from healthy controls were primarily naïve ^32^. In T1D, CD8^+^ T cells target and destroy insulin-producing β-cells in the pancreas. As GAD65 can be found in the brain but also in islets of Langerhans ^33,34^, a considerable number of patients with T1D have GAD65-antibodies in the serum and many patients with anti-GAD65 AINS do present with concomitant T1D ^4,35^. Previous studies identified an autoimmune stem-like CD8^+^ T cell population driving T1D ^33,36^. Those TSCM self-renew continuously and give rise to cytotoxic immune mediators destroying β-cells in the pancreas ^33^. Apart from T1D, stem-like T cells were found to maintain and promote the progression of other autoimmune diseases, e.g., systemic lupus erythematosus (SLE) and rheumatoid arthritis (RA). Frequencies of CD4^+^ TSCM cells were found to be significantly increased in the peripheral blood of SLE and RA patients when compared to healthy controls ^37,38^. Similar to these reports, we detected an expansion of activated CD4^+^ TSCM in the CSF of patients with anti-GAD65 AINS. We speculate that those CD4^+^ TSCM cells represent a long-lived reservoir of autoreactive T cells driving the immune response seen in anti-GAD65 AINS by continuously supporting CD8^+^ T cells and, thereby, maintaining and promoting the progression of autoimmune disease ^33,36,39^. CD4^+^ TSCM were especially increased in anti-GAD65 AINS patients with a disease duration over one year, whereas the increase in PB plasmablast was mainly noted in early disease stages. This further supports the concept of a persisting T cell-mediated immune response as described previously ^7,8,40^.

Apart from an increase in CD4^+^ TSCM, cluster analysis revealed higher proportions of MLC in the CSF of anti-GAD65 AINS compared to IIH patients. MLC are hematopoietic cells originating from the bone marrow. Under physiological conditions, the mature microglia population is mostly permanent and self-renews autonomously. In contrast, under pathological conditions, MLC can populate the CNS. In this context, an impaired blood-brain-barrier together with the release of several cytokines and chemokines as well as a reduction in CNS resident microglia contribute to this CNS engraftment. Despite their resemblance to microglia, MLC show different kinetics and functional profiles ^41^. However, our DGE analysis did not reveal any significant differences in gene expression of this cell population between anti-GAD65 AINS and IIH patients. Further studies are necessary to delineate the role of MLC in the pathogenesis of anti-GAD65 AINS and to assess their therapeutic potential.

Furthermore, DGE analysis revealed reduced expression of *HLA-C* in the PBMCs of anti-GAD65 AINS compared to IIH patients. *HLA-C* is a gene locus on chromosome 6 encoding HLA-C alleles, which are involved in the regulation of the innate and adaptive immune response ^42^. They can interact with NK cells (via their killer-cell immunoglobulin-like (KIR) receptors) and with CD8^+^ T cells as well as indirectly with CD4^+^ T cells ^42^. A recently published GWAS analysis of patients with anti-GAD65 AINS has identified *HLA-C* as one protein-coding gene mapping to the anti-GAD65 AINS GWAS loci ^9^. Likewise previous studies linked alterations in the *HLA-C* locus to other autoimmune diseases like psoriasis, Crohn’s disease, alopecia areata, and rheumatic diseases ^43–46^. Thus, *HLA-C* might be an interesting locus in the pathophysiology of anti-GAD65 AINS suggesting a need for more comprehensive research in that field.

Our analysis of TCR repertoires in anti-GAD65 AINS revealed distinct patterns of clonal expansion and compartmentalization between the CSF and PB. The observation of significantly expanded T cell clones in the CSF of anti-GAD65 AINS patients is consistent with findings from other autoimmune encephalitis studies, such as in anti-GABAa encephalitis, where Bracher et al. (2020) demonstrated that a CD8^+^ T cell clone was strongly expanded in the CSF ^47^. The reciprocal relationship between peripheral TCR diversity and CSF clonal expansion supports the notion of a focused autoimmune response, in which pathogenic T cells migrate from the periphery to the CNS, resulting in an accumulation of disease-relevant T cells within the CNS and leaving a less diverse T cell repertoire in the peripheral circulation. This pattern mirrors observations in MS, where Jacobsen et al. (2002) reported compartmentalized oligoclonal expansion of memory CD8^+^ T cells in the CSF ^30^. Moreover, Sousa et al. have also found that highly expanded T cell clones were enriched in the CSF compartment of MS patients compared to PB ^48^. The inverse relationship between CSF clonal expansion and peripheral diversity may represent a hallmark of CNS-directed autoimmunity, as similarly described in other inflammatory immune-mediated diseases of the CNS ^49^. The distinct clonal network patterns in anti-GAD65 AINS patients, characterized by larger interconnected clusters of expanded TSCM cells and the high clonal frequencies observed in the top 20 most abundant TCR clones of anti-GAD65 AINS patients’ CSF TSCM further support the notion of a focused, disease-relevant immune response. This is consistent with the concept of epitope-spreading in autoimmune diseases, where the initial autoimmune response against a specific antigen (in this case, GAD65) leads to the activation and expansion of T cells recognizing additional epitopes. Future research should focus on characterizing the antigen specificity of these expanded clones and exploring their potential as therapeutic targets or biomarkers for disease progression and treatment response. Notably, our finding of distinct transcriptional profiles in expanded CSF clones, particularly the upregulation of T cell proliferation and inflammatory response genes, is consistent with observations in other neuroinflammatory conditions like MS, which was reported by Pappalardo et al. (2020) ^50^. Moreover, a study by Gate et al. on Alzheimer’s disease found that clonally expanded CD8^+^ T cells in the CSF showed increased expression of cytotoxic effector genes, including *NKG7* and *GZMH* ^51^. This finding suggests that the upregulation of activation- and inflammatory response-related genes in expanded CSF clones may be a common feature across various neurodegenerative and autoimmune conditions affecting the CNS.

In contrast to the clonal expansion patterns observed in the TCR repertoire, we did not identify relevant clonally expanded B cells in the CSF of anti-GAD65 AINS patients. However, recombinant production and reactivity testing of mAbs derived from randomly chosen CSF BCRs with concurrently available heavy and light chains revealed a remarkably high frequency of GAD65-reactive mAbs. Somatic hypermutations were increased in GAD65-reactive as compared to non-reactive mAbs, suggesting that they likely have undergone antigen-driven affinity maturation, as described previously ^52^. In contrast to a previous study ^52^ that identified two (out of seven) patients with GAD65-reactive BCRs derived from antibody secreting cells (ASCs) in the CSF, we did not find GAD65-reactive BCRs in CSF ASCs of the four patients included in our BCR analysis, indicating that only few patients harbor them. As Tröscher et al. ^8^ have found plasma cell infiltrations into the brain parenchyma in GAD-TLE patients at early disease stages, another point to consider is that GAD65-reactive plasma cells could have already evaded the CSF by the time of sampling in our study. Similarly, the increase in plasmablasts in the PB was mainly found in patients with a disease duration below one year in our anti-GAD65 AINS cohort, suggesting that the B cellular immune response in anti-GAD65 AINS patients might be restricted to early disease stages.

Our study is limited by the small sample size and the differences in age and sex between anti-GAD65 AINS and IIH patients. Different disease durations within the anti-GAD65 AINS cohort further limit our analyses. In addition, different kits were used for library preparation.

However, our study exceeds previous ones by providing an in-depth characterization of peripheral and intrathecal immune cell populations in mainly treatment-naive anti-GAD65 AINS patients compared to non-inflammatory controls, yielding new insights into disease pathogenesis and identifying potential novel treatment targets.

In conclusion, our findings contribute to the growing body of evidence supporting the role of antigen-specific B and T cell responses in autoimmune encephalitis. The compartment-specific changes in the TCR repertoire and clonal expansion patterns emphasize the importance of studying both CSF and PB to improve the understanding of the pathogenesis of CNS autoimmune disorders. Our findings highlight the potential role of CD4^+^ TSCM cells in the pathogenesis of anti-GAD65 AINS and suggest that these cells may serve as a reservoir for disease-specific T cells in the CNS. Future research should focus on characterizing the antigen specificity of expanded CD4^+^ TSCM clones, clarifying T-B-cell interactions, and assessing their potential as therapeutic targets or biomarkers for disease progression and treatment responses.

## STAR Methods

### Experimental model and study participant details

Patients were prospectively included in the GAD sequencing-cohort at four specialized clinical centers (Münster, Rotterdam, Kiel, and Munich) according to the following inclusion criteria:

1. Suspected anti-GAD65 AINS presenting with one of the typical clinical phenotypes or confirmed anti-GAD65 AINS.
2. Performance of a lumbar puncture (LP) during clinical routine workup.
3. No ongoing or previous immunomodulatory therapy (treatment-naïve) except from IVIG, steroids, immunoadsorption or plasmapheresis at least 6 months prior to sampling.
4. Written informed consent to participate in the study.

Patients were excluded if one or more of the following exclusion criteria applied:

1. Questionable diagnosis.
2. Signs of acute infection, assessed by routine blood and CSF analysis.
3. Known comorbid systemic autoimmune disease (e.g., systemic lupus erythematosus) or chronic infections (e.g. HIV, chronic hepatitis), except for T1D.
4. Pregnancy or breastfeeding.
5. Age < 18 years at time of inclusion.
6. Mental illness impairing the ability to give informed consent.
7. Artificial blood contamination of the CSF resulting in >200 erythrocytes/μL. The study design is illustrated in **Figure 1A**.

Anti-GAD65 AINS patients included in the final study cohort had high IgG-autoantibodies (AABs) against GAD65 in serum as defined by the presence of a typical staining pattern on rodent or non-human primate brain tissue and pancreas. In addition, GAD65-AABs could be detected in the CSF by immunohistochemistry on rodent or non-human primate brain tissue and cell-based assays. In one patient, GAD65-AABs were only analyzed by Enzyme-linked Immunosorbent Assay (ELISA).

Due to ethical issues to perform a LP in healthy controls, patients with idiopathic intracranial hypertension (IIH) served as non-inflammatory controls (n = 8) ^53^ ^54^ (**Figure 1A**). All patients met the inclusion criteria 3 & 4 and none of the above-mentioned exclusion criteria. Parts of the IIH cohort have been published before ^54,55^. Detailed clinical data of the anti-GAD65 AINS and IIH cohorts are summarized in **Supplementary Table 1-4.**

Samples were processed according to a standardized protocol at all centers minimizing the risk for systematic technical bias. The study was performed in accordance with the declaration of Helsinki and was approved by the local ethics committees: (1) the “Ethikkommission der Ärztekammer Westfalen-Lippe (ÄKWL) und der Westfälischen-Wilhelms-Universität” (Ethics Committee of the Board of Physicians of the Region Westfalen-Lippe and of the Westfälische Wilhelms-University; reference number 2015-522-f-S) in Münster, (2) the “Erasmus MC Medical Ethical Research Board” (reference numbers: MEC-2015-397, MEC-2020-0418, and MEC-2020-0650) in Rotterdam, (3) the “Ethikkommission” of the Medical Faculty Kiel (reference numbers: D498/19, B337/13, B300/19, and D578/18), and (4) by the “Ethikkommission” of the Medical Faculty Munich (projects: 18-419, 163-16, 22-0150, 173-14).

### Method details

#### Collection of PB and CSF samples

PB and CSF samples were collected as previously described ^53^. In brief, up to 30 mL of CSF and 3 mL of PB were collected during clinical routine workup, in addition to the diagnostic samples. To ensure optimal sample quality, CSF was transported at 4°C and samples were processed within one hour of sample collection. Basic CSF parameters (CSF cell count, concentration of protein, albumin, immunoglobulins [Ig] in serum and CSF as well as OCBs) were analyzed as part of the clinical routine workup according to the standard diagnostic procedures of each center. A Reiber scheme was used to assess the integrity of the blood-CSF barrier (BCSFB). CSF was centrifuged at 300×g for 10 min and the supernatant was removed.

For scRNA-seq, CSF cells were resuspended in 5 mL of X-Vivo15 media (Lonza). 5 μl of the single-cell suspension were manually counted using a counting chamber. Up to 20,000 CSF cells were loaded onto the 10x Genomics chip for scRNA-seq.

#### Single cell RNA-sequencing

Samples were analyzed in an unbiased fashion using scRNA-seq as described previously ^53,55^. In brief, using the Chromium Single Cell GEM, single-cell suspensions were loaded onto the Chromium Single Cell Controller Library & Gel Bead Kits (10X Genomics). For 5’ reagents, chemistry version V1 and V1.1 and for 3’ reagents, V3 and V3.1 were used. Samples and libraries were prepared according to the manufacturer’s instructions. Sequencing was performed on an Illumina NextSeq500 or an Illumina Novaseq 6000 with a 2 x 150 or 2 x 100 read setup.

#### BCR cloning, recombinant mAb production and anti-GAD65 reactivity screening

Corresponding full length consensus sequences of variable heavy (VH) and light chains (VL) of a representative amount of CSF B cells were retrieved from VDJ Cell Ranger data. Sequences were synthesized by GeneArt (Thermo Fisher) and cloned into the expression vector pTT5. Ig vector pairs were transiently transfected into HEK EBNA cells, and mAb purification was performed by immobilized metal affinity chromatography, as described previously ^52,56^. Antibody concentrations were determined by BCA (Thermo Fisher) and mAb integrity was verified by SDS-PAGE and Coomassie blue staining (Thermo Fisher). GAD65-reactivity was tested by both cell-based-assay (Euroimmun) and indirect immunofluorescence staining on primate brain slides (Euroimmun), according to the manufacturer’s instructions, respectively.

### Quantification and statistical analysis

#### Bioinformatic analyses of single cell RNA-sequencing data

The 10x Genomics’ Cell Ranger (v6.1.2) multi pipeline with the transcriptome reference (refdata-gex-GRCh38-2020-A) and V(D)J reference (refdata-cellranger-vdj-GRCh38-alts-ensembl-7.1.0) was used for data processing.

Filtered feature-barcode matrix files were then analyzed with the R package ‘Seurat (v5.0.1)’^57^. Initially, quality control was performed to ensure the reliability of the data. This involved analyzing and visualizing the number of RNA features per cell, the proportion of mitochondrial RNA, and ribosomal RNA for each cell. QC thresholds for nFeature RNA, nCount RNA, mitochondrial/ribosomal gene percentages, and log10GenePerUMI were determined using sample-specific outlier detection (using isOutlier() function from the scuttle package) and validated through violin plots. FeatureScatter visualizations (nCount RNA vs. nFeature RNA, nCount RNA vs. mitochondrial%, nCount RNA vs. ribosomal%) further guided threshold selection by revealing relationships between RNA content, technical artifacts, and biological states (https://doi.org/10.1093/bioinformatics/btw777). Cells with a low or very high number of RNA features and/or a high amount of mitochondrial/ribosomal RNA were removed from the dataset as those present low-quality cells or doublets. In addition, doublet removal was performed with the R package ‘DoubletFinder’ (v2.0.3) ^58^. Artificial doublets are computationally generated by merging transcriptomic profiles from the datasets. The combined real and synthetic data undergo preprocessing, including normalization and dimensionality reduction, to ensure compatibility. Principal component analysis (PCA) is applied, followed by k-nearest neighbor analysis in the reduced space to calculate each cell’s proportion of artificial neighbors (pANN). Cells are then ranked by pANN scores and classified as doublets using a threshold based on the expected doublet frequency in the experiment. Next, normalization was performed using the function ‘SCTransform’ ^59^. Variable features were identified, and features were scaled and centered in the dataset. The RunPCA function was used to perform a principal component analysis (PCA) to reduce the dimensionality of the dataset (20 dimensions). Data integration was performed with the Seurat function IntegrateLayers() using method = HarmonyIntegration ^60^ to correct for effects caused by different library preparation kits (3’ vs 5’) and different sampling locations ^60^. A shared nearest neighbor graph was created with the FindNeighbors function incorporating the first 20 principal components, and clusters were identified with the FindClusters function using a resolution of 0.5. This optimal number of dimensions was determined through an automated process that evaluates the information content of higher-order PCs. Finally, the Uniform Manifold Approximation and Projection (UMAP) dimensional reduction was applied using the RunUMAP function with 20 dimensions. The clusters were annotated using a combination of manual curation and automated methods with the ‘Azimuth’ package (v0.5.0), leveraging the human peripheral blood mononuclear cells (PBMC) reference dataset ^57^. For further confirmation, the clusters were automatically annotated with the R packages ‘SingleR’ (v2.4.0) based on the Immune Cell Expression reference dataset ^61^. Four clusters were identified as low-quality cells based on the expression of different mitochondrial genes and/or inconsistent marker gene expression. Consequently, these clusters were removed from the Seurat object.

#### Cell abundance analysis and cluster-specific differential gene expression

The R package ‘propeller’ (part of speckle v1.2.0) was used to assess differentially abundant clusters across all cell types between anti-GAD65 AINS and IIH patients^62^. Data were log-transformed and Wilcoxon signed-rank test was performed. Volcano plots of cluster abundances were created with ggplot2 (v3.4.4) ^63^.

To perform cluster-specific differential gene expression, first pseudobulking was performed using the function ‘AggregateExpression’. Next, the ‘FindMarkers’ (DESeq2) was applied to every cluster to compare their gene expression between anti-GAD65 AINS and IIH patients. Differential gene expression analysis comparing *expanded* and *non-expanded T-cell receptor (TCR) clonotypes* was conducted using TCR-annotated single-cell RNA sequencing data. PyDESeq2, a negative binomial model-based framework was used to identify transcriptional signatures associated with clonal expansion dynamics ^64^. The R package ‘EnhancedVolcano’ (v1.20.0) ^65^ was used to create volcano plots of differentially expressed genes. All analyses were performed separately for PBMCs and CSF cells. Gene Ontology (GO) enrichment analysis of differentially expressed genes from anti-GAD65 AINS patients was performed using R based ‘clusterprofiler’ (v4.16.0) (doi: 10.1016/j.xinn.2021.100141). The R package org.Hs.eg.db (v3.19) was used for Genome wide annotation for Human.

#### BCR and TCR bioinformatic analysis

Single-cell immune receptor repertoire analysis was performed on Cell Ranger Multi pipeline-processed data using the Python-based Dandelion package (v0.3.8) ^66^. Cellranger multi pipeline processed VDJ sequences were aligned against the IgBLAST database and germline sequences using the Dandelion ^66^ package. The analysis pipeline integrated both Dandelion pre-processed VDJ output files and Seurat-converted transcriptomic AnnData objects using a combination of the Dandelion, Scanpy (v1.10.2) ^67^, and Scirpy (v0.17.2) ^68^ packages. For TCR analysis, we selected cells containing TRB chains paired with their corresponding TRA chains, including additional VJ loci. Similarly, for BCR analysis, we retained cells with IGH chains paired with their corresponding IGL chains and additional VJ loci. Clonal relationships were visualized through network analysis, and clonotype distributions were quantified using clonal frequency plots. Comprehensive clonotype characterization, including diversity metrics and repertoire analysis, was conducted using the Dandelion and Scirpy computational frameworks and Python- and R-based custom in-house pipelines. Somatic hypermutation (SHM) frequencies in BCR sequences and GAD65-reactive antibody sequences were quantified using the Change-O and SHazaM framework within the Immcantation portal, enabling comprehensive analysis of B cell affinity maturation patterns ^69^.

Based on the Cell Ranger output, the Gini-Simpson diversity was determined with the diversity function of the R microbiome package v1.23.1 ^70^, while the Shannon and the inverse simpson index were calculated with the diversity function of the R vegan package v2.6.4 ^71^. The reported cell frequencies for the clones were visualized in the UMAP plots. A clone in the context of BCR or TCR repertoire analysis is defined by a set of specific criteria demonstrated at Dandelion documentation ^66^. Firstly, all members of a clone must share identical V- and J-gene usage in their VDJ chain, which applies to IGH for B cells or TRB/TRD for T cells. Secondly, the CDR3 junctional or CDR3 sequence length in the VDJ chain must be identical across all clonal members. Thirdly, the VDJ chain junctional or CDR3 sequences must meet a minimum sequence similarity threshold, typically calculated using the Hamming distance. This similarity cut-off is adjustable, with a default of 85% for BCRs, but it should be set to 100% when analyzing TCR data due to their more conserved nature ^66^. Lastly, the algorithm considers VJ chain usage (IGK/IGL for B cells, TRA/TRG for T cells). If cells within a putative clone exhibit different VJ chains, the clone is further split following the same conditions applied to VDJ chains. To determine the somatic hypermutation, clones with at least three cells were extracted and the consensus sequence of each clone aligned with all human IGHV reference sequences annotated in the IMGT database ^72^. The reference sequence with the highest identity was then compared with all sequences of the cells and the mean number of mutations per clone was determined. In addition, we performed for each clone a hierarchical clustering using the R function hclust out of the stats package v4.2.2 ^73^. TCR sequence-sharing was assessed with GLIPH v2 ^74,75^ (73, 74) and TCR antigen association with VDJdb ^76^.

### Key resource table

| REAGENT or RESOURCE | SOURCE | IDENTIFIER |
| --- | --- | --- |
| **Biological samples** |  |  |
| Cerebrospinal fluid (CSF) | anti-GAD65 AINS and IIH patients | NA |
| Peripheral Blood Mononuclear Cells (PBMCs) | anti-GAD65 AINS and IIH patients | NA |
| **Chemicals, peptides, and recombinant proteins** |  |  |
| X-Vivo15 media | Lonza | #02-053Q |
| T4 Polynucleotide Kinase | New England Biolabs | #M0201S |
| T4 DNA Ligase | New England Biolabs | #M0202S |
| RecJF | New England Biolabs | #M0264S |
| Lambda Exonuclease | New England Biolabs | #M0262S |
| Freestyle medium | Thermo Fisher Scientific | 12338-018 |
| 10% Pluronic F68 | Thermo Fisher Scientific | 24040-032 |
| Geneticin | Thermo Fisher Scientific | 10131-035 |
| Phusion High-Fidelity DNA Polymerase | New England Biolabs | M0530L |
| BssHII | New England Biolabs | R0199S |
| Sall-HF | New England Biolabs | R3138S |
| KasI | New England Biolabs | R0544S |
| SacII | New England Biolabs | R0157S |
| dNTPs | Thermo Fisher Scientific | R0181 |
| Pierce Coomassie Brilliant Blue Dye | Thermo Fisher Scientific | 20278 |
| PEI | Polysciences | 23966-2 |
| **Critical commercial assays** |  |  |
| Chromium Next GEM Single Cell 3' Library & Gel Bead Kit v3 | 10x Genomics | PN-1000075 |
| Chromium Next GEM Single Cell 3' Library & Gel Bead Kit v3.1 | 10x Genomics | PN-1000121 |
| Chromium Next GEM Single Cell 5' Library & Gel Bead Kit v1 | 10x Genomics | PN-1000156 |
| Chromium Next GEM Single Cell 5' Library & Gel Bead Kit v1.1 | 10x Genomics | PN-1000165 |
| Chromium Single Cell VDJ Enrichment Kit, Human T Cell | 10x Genomics | PN-1000005 |
| Chromium Single Cell VDJ Enrichment Kit, Human B Cell | 10x Genomics | PN-1000016 |
| Glutamatdecarboxylase (GAD) 65 kDa IIFT | Euroimmun | FA 1022-1005-50 |
| Neurology Mosaic 17 | Euroimmun | FB 1111-1010-17 |
| Ni-NTA Spin Columns | Qiagen | 31014 |
| D-Tube Dialyzer Maxi, MWCO 12-14 kDa | Merck | 71510-3 |
| Pierce BCA Protein Assay Kit | Thermo Fisher Scientific | 23227 |
| **Deposited data** |  |  |
| Single-cell sequencing and clinical data | ABCD-J data catalog | https://data.abcd-j.de/dataset/d1b56ad8-f8e9-5d4b-aa76-fe1c155ad201/1.0 |
| **Experimental models: Cell lines** |  |  |
| HEK EBNA cells | supplied by Dr. Jenne (Perera et al., 2013) | NA |

| **Experimental models: Organisms/strains** |  |  |
| --- | --- | --- |
| 8 patients with GAD65-antibody associated autoimmune neurological syndromes (anti-GAD65 AINS) | specialized clinical centers (Münster, Rotterdam, Kiel, and Munich) | NA |
| 8 patients with Idiopathic Intracranial Hypertension (IIH) | specialized clinical centers (Münster, Rotterdam, Kiel, and Munich) | NA |
| **Software and algorithms** |  |  |
| Cell Ranger (v6.1.2) multi pipeline | 10x Genomics | https://www.10xgenomics.com/support/software/cell-ranger/latest |
| Cell Ranger software v3.1.0 | 10x Genomics | https://www.10xgenomics.com/support/software/cell-ranger/latest |
| propeller (part of speckle v1.2.0) | Belinda Phipson | https://doi.org/10.1093/bioinformatics/btac582 |
| ggplot2 (v3.4.4) | Hadley Wickham | https://doi.org/10.1007/978-3-319-24277-4 |
| EnhancedVolcano (v1.20.0) |  |  |
| Scanpy (v1.10.2) | F. Alexander Wolf | https://doi.org/10.1186/s13059-017-1382-0 |
| Scirpy (v0.17.2) | Gregor Sturm | https://doi.org/10.1186/s13059-017-1382-0 |
| Change-O and SHazaM framework within the Immcantation portal | Namita T. Gupta | https://doi.org/10.1093/bioinformatics/btv359 |
| VDJ reference GRCh38 v3.1.0 | 10x Genomics | https://www.10xgenomics.com/support/software/cell-ranger/latest/release-notes/cr-reference-release-notes |
| microbiome R package (v1.23.1) | Leo Lahti, Sudarshan Shetty | http://microbiome.github.com/microbiome |
| vegan package (v2.6.4) | Jari Oksanen | https://github.com/vegandevs/vegan |
| stats package (v4.2.2) | R Foundation for Statistical Computing | https://www.R-project.org/ |
| GLIPH v2 | Huang Huang, Jacob Glanville | https://doi.org/10.1038/s41587-020-0505-4<br>https://doi.org/10.1038/nature22976 |
| VDJdb | Mikhail Shugay | https://doi.org/10.1093/nar/gkx760 |
| IMGT database |  | [10.1101/pdb.top115](https://doi.org/10.1101/pdb.top115). |
| Dandelion package (v0.3.8) | Zewen Kelvin Tuong | https://doi.org/10.1038/s41587-023-01734-7 |
| **Other** |  |  |
| pTT5 (IgG1 expression vector) | supplied by Dr. Jenne (Perera et al., 2013) | NA |
| GeneArt Gene Synthesis | Thermo Fisher Scientific | NA |

## Data and code availability

Data underlying this study are registered with the ABCD-J data catalog at https://data.abcd-j.de/dataset/d1b56ad8-f8e9-5d4b-aa76-fe1c155ad201/1.0. Regarding the computational analyses, the official tutorial of the packages listed were followed and a custom in-house pipeline was used. The codes used in this study are available from the corresponding author upon reasonable request.

## Acknowledgements

This project was supported by: the German Research Foundation (ERARE18-202 UltraAIE) and the Netherlands Medical Research Foundation (ZonMW) (ERARE-JTC2018 202: UltraAIE) under the frame of E-Rare-3, the ERA-Net for Research on Rare Diseases; the German Federal Ministry of Education and Research (Comprehensive, Orchestrated, National Network to Explain, Categorize and Treat autoimmune encephalitis and allied diseases within the German NEtwork for Research on AuToimmune Encephalitis – CONNECT GENERATE and CONNECT GENERATE 2.0; 01GM1908A and 01GM2208A); Dioraphte (2001 0403); EpilepsieNL (19-18); the Research Committee of the Faculty of Medicine of the Heinrich Heine University Düsseldorf (grant number 2022-04); the ‘Else Kröner-Fresenius Stiftung’ (grant number 2023_EKMS.05); the Fritz Thyssen Stiftung (grant number 10.23.1.015MN); LMU excellent (grant number: AOST 867603-4); and the Friedrich-Baur-Stiftung (54/23).

We would like to thank all participating patients and their relatives and caregivers for their invaluable contribution to this study. We acknowledge High-Performance Computing (HPC) support from the Centre for Information and Media Technology (ZIM) at the Heinrich Heine University Düsseldorf. This work was part of Saskia Räuber’s and Paul Disse’s PhD research.

## Author Contributions

**SB**: Methodology, Software, Validation, Formal analysis, Data Curation, Writing – original draft preparation, Writing – review and editing, Visualization; **SR:** Methodology, Software, Validation, Formal analysis, Investigation, Data Curation, Writing – original draft preparation, Writing – review and editing, Visualization, Resources, Funding acquisition; **KE**: Methodology, Software, Validation, Formal analysis, Investigation, Data Curation, Writing – original draft preparation, Writing – review and editing, Resources, Visualization, Funding acquisition; **DE**: Formal analysis, Software, Writing - review and editing; **MvD**: Software, Writing - review and editing; **MS**: Methodology, Validation, Formal analysis, Writing - review and editing; **MH**: Resources, Writing - review and editing; **PD**: Formal analysis, Writing - review and editing; **MJ**: Resources; **DP**: Resources; **MH**: Software, Investigation, Resources, Writing - review and editing; **LMM**: Investigation, Resources,Writing - review and editing; **MP**: Resources; **CS**: Resources; **EH**: Investigation; **EK**: Resources, Writing - review and editing; **JD**: Investigation, Resources; **SK**: Resources; **MR**: Resources; **EB**: Resources; **HW**: Resources; **NHS**: Resources; **JF**: Resources; **NG**: Resources; **LK**: Resources; **MR**: Resources, Writing - review and editing; **AR**: Resources,Writing - review and editing; **MS**: Resources; **SGM**: Resources; **MJT**: Investigation, Resources, Supervision, Funding acquisition, Project administration, Writing – review and editing; **FL**: Investigation, Resources, Supervision, Funding acquisition, Project administration, Writing – review and editing; **GMzH**: Investigation, Resources, Supervision, Funding acquisition, Project administration, Writing – review and editing; **FT**: Investigation, Resources, Supervision, Project administration, Writing – review and editing, Funding acquisition; **NM**: Investigation, Resources, Supervision, Funding acquisition, Project administration, Writing – review and editing.

## Consortia

**The EMC AIE study group**: Juna M de Vries, Mariska M. P. Nagtzaam, Suzanne C. Franken, Yvette S. Crijnen, Juliette Brenner, Robin W. van Steenhoven, Jeroen Kerstens, Marienke A. A. M. de Bruijn, Anna E. M. Bastiaansen, Remco M. Hoogenboezem, Sharon Veenbergen, Peter A. E. Sillevis Smitt.

**The GENERATE study group:** Michael Adelmann, Luise Appeltshauser, Ilya Ayzenberg, Carolin Baade-Büttner, Andreas van Baalen, Sebastian Baatz, Oliver Bähr, Bettina Balint, Sebastian Bauer, Annette Baumgartner, Stefanie Becker, Sonka Benesch, Robert Berger, Birgit Berger, Martin Berghoff, Sascha Berning, Sarah Bernsen, Achim Berthele, Christian Bien, Corinna Bien, Andreas Binder, Stefan Bittner, Daniel Bittner, Franz Blaes, Astrid Blaschek, Amelie Bohn, Sergio Castro-Gomez, Justina Dargvainiene, Timo Deba, Julia Maren Decker, Andre Dik, Juliane Dominik, Kathrin Doppler, Mona Dreesmann, Friedrich Ebinger, Lena Edelhoff, Laura Ehrhardt, Sven Ehrlich, Katharina Eisenhut, Alexander Emmer, Dominique Endres, Marina Entscheva, Daniela Esser, Thorleif Etgen, Jürgen Hartmut Faiss, Kim Kristin Falk, Walid Fazeli, Alexander Finke, Carsten Finke, Dirk Fitzner, Marina Flotats-Bastardas, Mathias Fousse, Tobias Freilinger, Paul Friedemann, Manuel Friese, Marco Gallus, Marcel Gebhard, Christian Geis, Anna Gorsler, Armin Grau, Oliver Grauer, Britta Greshake, Catharina Groß, Thomas Grüter, Aiden Haghikia, Robert Handreka, Niels Hansen, Jens Harmel, Antonia Harms, Yetzenia Dubraska Haro Alizo, Martin Häusler, Joachim Havla, Chung Ha-Yeun, Wolfgang Heide, Valentin Held, Kerstin Hellwig, Philip Hillebrand, Frank Hoffmann, Christian Hofmann, Ulrich Hofstadt-van Oy, Peter Huppke, Hagen Huttner, Fatme Seval Ismail, Martina Jansen, Mareike Jansen, Aleksandra Juranek, Michael Karenfort, Max Kaufmann, Christoph Kellinghaus, Constanze Kerin (geb. Mönig), Jaqueline Klausewitz, Susanne Knake, Ellen Knierim, Peter Körtvélyessy, Stjepana Kovac, Andrea Kraft, Markus Krämer, Verena Kraus, Christos Krogias, Gregor Kuhlenbäumer, Tanja Kümpfel, Christoph Lehrich, Jan Lewerenz, Frank Leypoldt, Andeas Linsa, Jan Lünemann, Marie Madlener, Michael Malter, Niels Margraf, Carlos Martinez Quesada, Monika Meister, Nico Melzer, Kristin Stefanie Melzer, Til Menge, Sven Meuth, Gerd Meyer zu Hörste, Fabian Möller, Marie-Luise Mono, Sigrid Mues, Jost Obrocki, Loana Penner, Lena Kristina Pfeffer, Thomas Pfefferkorn, Steffen Pfeuffer, Alexandra Philipsen, Johannes Piepgras, Felix von Poderwils, Mosche Pompsch, Josef Priller, Anne-Katrin Pröbstel, Harald Prüß, Daniel Rapp, Dominica Ratuszny, Johanna Maria Helena Rau, Saskia Räuber, Robert Rehmann, Ina Reichen, Gernot Reimann, Raphael Reinecke, Nele Retzlaff, Marius Ringelstein, Henrik Rohner, Felix Rosenow, Kevin Rostasy, Theodor Rüber, Stephan Rüegg, Yannic Saathoff, Jens Schaumberg, Ruth Schilling, Mareike Schimmel, Jens Schmidt, Ina-Isabelle Schmütz, Hauke Schneider, Patrick Schramm, Stephan Schreiber, Gesa Schreyer, Ina Schröder, Simon Schuster, Günter Seidel, Frank Seifert, Thomas Seifert-Held, Makbule Senel, Kai Siebenbrodt, Olga Simova, Claudia Sommer, Juliane Spiegler, Oliver Stammel, Andeas Steinbrecher, Henning Stolze, Muriel Stoppe, Karin van’s Gravesande Storm, Christine Strippel, Dietrich Sturm, Klarissa Hanja Stürner, Kurt-Wolfram Sühs, Steffen Syrbe, Pawel Tacik, Simone Tauber, Franziska Thaler, Florian Then Bergh, Anja Tietz, Corinna Trebst, George Trendelenburg, Regina Trollmann, Thanos Tsaktanis, Hayrettin Tumani, Methap Türedi, Christian Urbanek, Niklas Vogel, Max Vogtmann, Matthias von Mering, Judith Wagner, Jan Wagner, Klaus-Peter Wandinger, Robert Weissert, Jonathan Wickel, Brigitte Wildemann, Karsten Witt, Kartharina Wurdack & Lara Zieger.

