## Supplementary Figure 1 for "CSF single-cell RNA sequencing reveals clonally expanded CD4^+^ stem cell-like memory T cells in GAD65-antibody associated neurological syndromes"

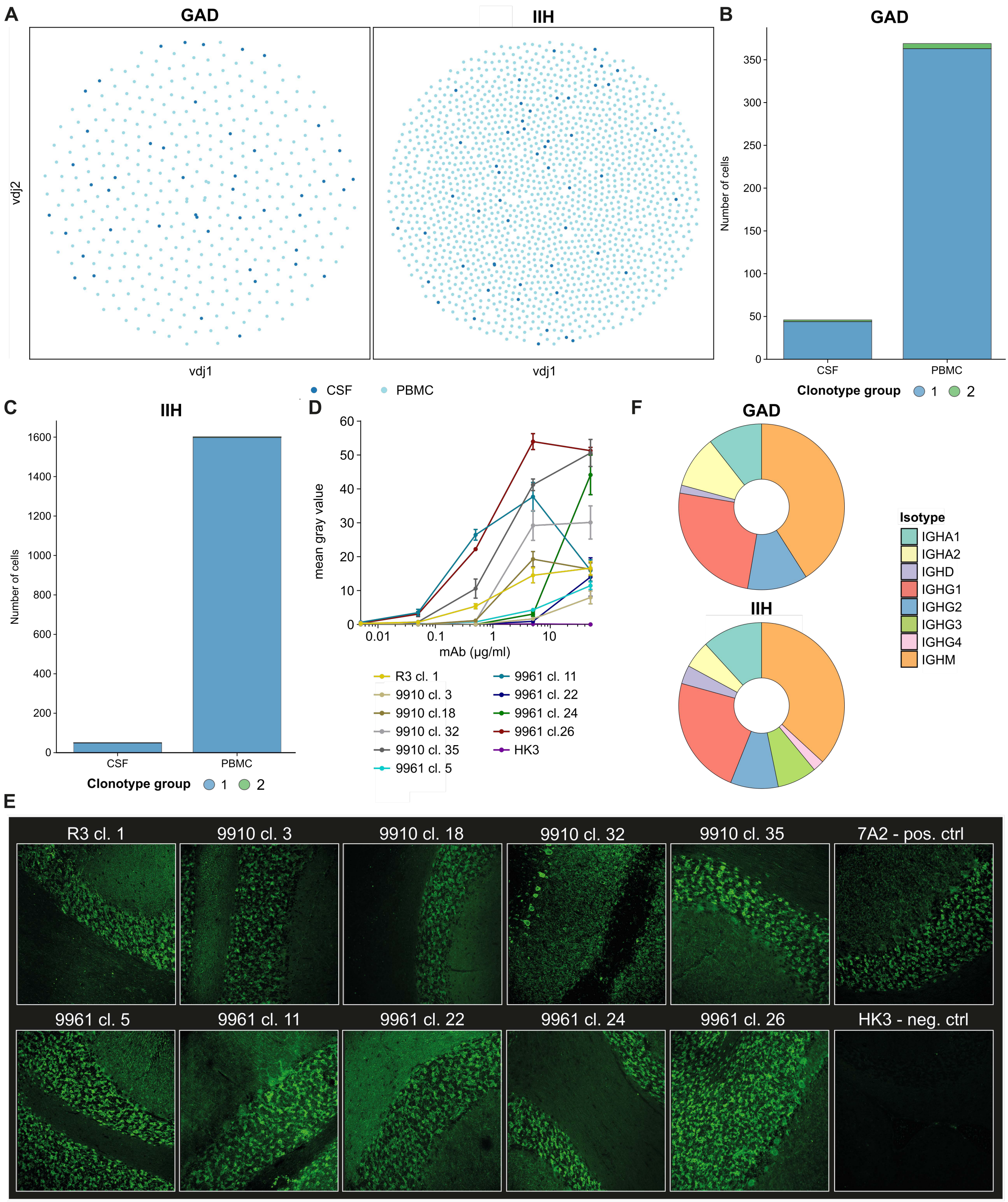


**Supplementary Figure 1 - Immunoglobulin isotype distribution of CSF BCRs, recombinant monoclonal antibody titration and primate tissue stainings.**

**A** Clonal network visualization depicting BCR repertoire relationships between CSF and PBMC compartments in anti-GAD65 AINS and IIH patients. Node size indicates clonal frequency. **B** Stacked bar plot showing quantitative comparison of BCR clonal distribution in CSF and PBMC compartments of anti-GAD65 AINS and IIH patients. Each stack shows the relative frequency of a distinct clonal group. **D** Relative binding strengths of the 10 GAD65-reactive mAbs were quantified by mAb serial dilutions and stainings of GAD65-transfected HEK cells. Mean gray values are shown as mean ± SEM. **E** Representative images of mAb stainings (50 µg/ml) on non-human primate cerebellar tissue. 7A2 serves as positive and HK3 as negative control. Stainings of all 10 GAD65-reactive mAbs are shown (as indicated by patient ID and clonotype number in column caption), all displaying a leopard-like staining of the cerebellar granular layer, typical for GAD65-specific tissue-reactivity. 9910 cl. 32, 9961 cl. 11 and cl. 26 additionally show some Purkinje-cell reactivity. **F** Relative immunoglobulin subclass distribution of all CSF BCRs in anti-GAD65 AINS and IIH patients.

*cl. - clonotype; CSF - cerebrospinal fluid; GAD - patients with glutamic acid decarboxylase antibody-associated autoimmune neurological syndromes (AINS); mAb - monoclonal antibody; neg. ctrl - negative control; PBMC - Peripheral Blood Mononuclear Cell, pos. ctrl - positive control*
