## Supplementary Figure 2 for "CSF single-cell RNA sequencing reveals clonally expanded CD4^+^ stem cell-like memory T cells in GAD65-antibody associated neurological syndromes"

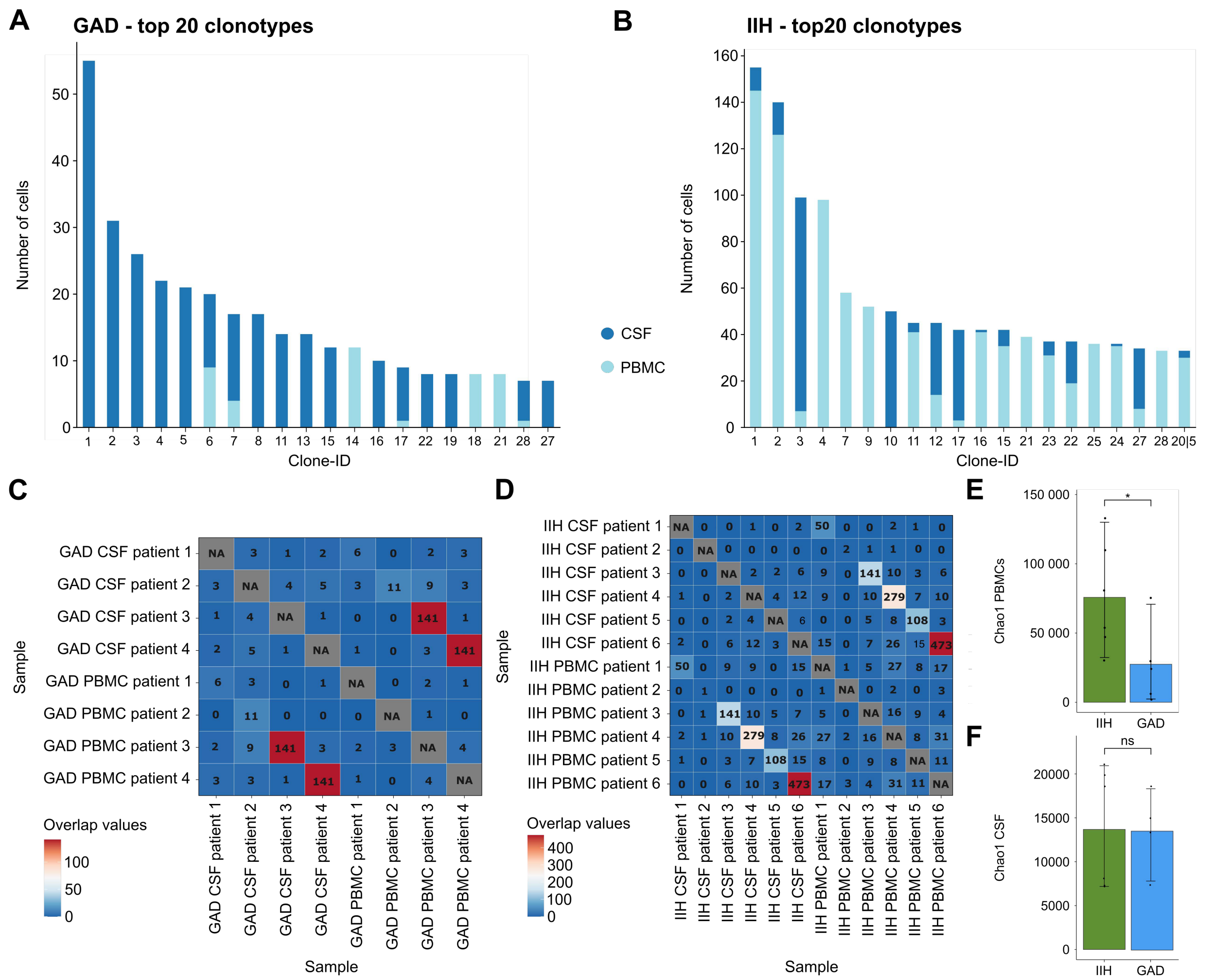


**Supplementary Figure 2 - The top clonotype composition of T cell repertoire in anti-GAD65 AINS and IIH patients.**

**A** Bar plot represents comparative analysis of the top 20 most abundant TCR clones in anti-GAD65 AINS patients, showing relative frequencies between CSF and PBMC compartments. **B** Bar plot represents comparative analysis of the top 20 most abundant TCR clones in IIH patients, showing relative frequencies between CSF and PBMC compartments. **C** Repertoire overlap heatmap of anti-GAD65 AINS patients represents repertoire similarity between CSF and PBMC compartment and among patients. **D** Repertoire overlap heatmap of IIH patients represents repertoire similarity between CSF and PBMC compartment and among controls. **E** TCR diversity metrics in the PBMC compartment, comparing anti-GAD65 AINS and IIH patients. Data shown as Chao1 index with statistical significance indicated. Statistical significance indicated where applicable (*p<0.05). **F** TCR diversity metrics in the CSF compartment, comparing anti-GAD65 AINS and IIH patients. Data shown as Chao1 index with no statistical significance indicated.

*GAD - patients with glutamic acid decarboxylase antibody-associated autoimmune neurological syndromes (AINS); CSF - cerebrospinal fluid; ns - not significant; PBMC - Peripheral Blood Mononuclear Cell; IIH - idiopathic intracranial hypertension; TCR - T cell receptor.*
