## Supplementary Figure 3 for "CSF single-cell RNA sequencing reveals clonally expanded CD4^+^ stem cell-like memory T cells in GAD65-antibody associated neurological syndromes"

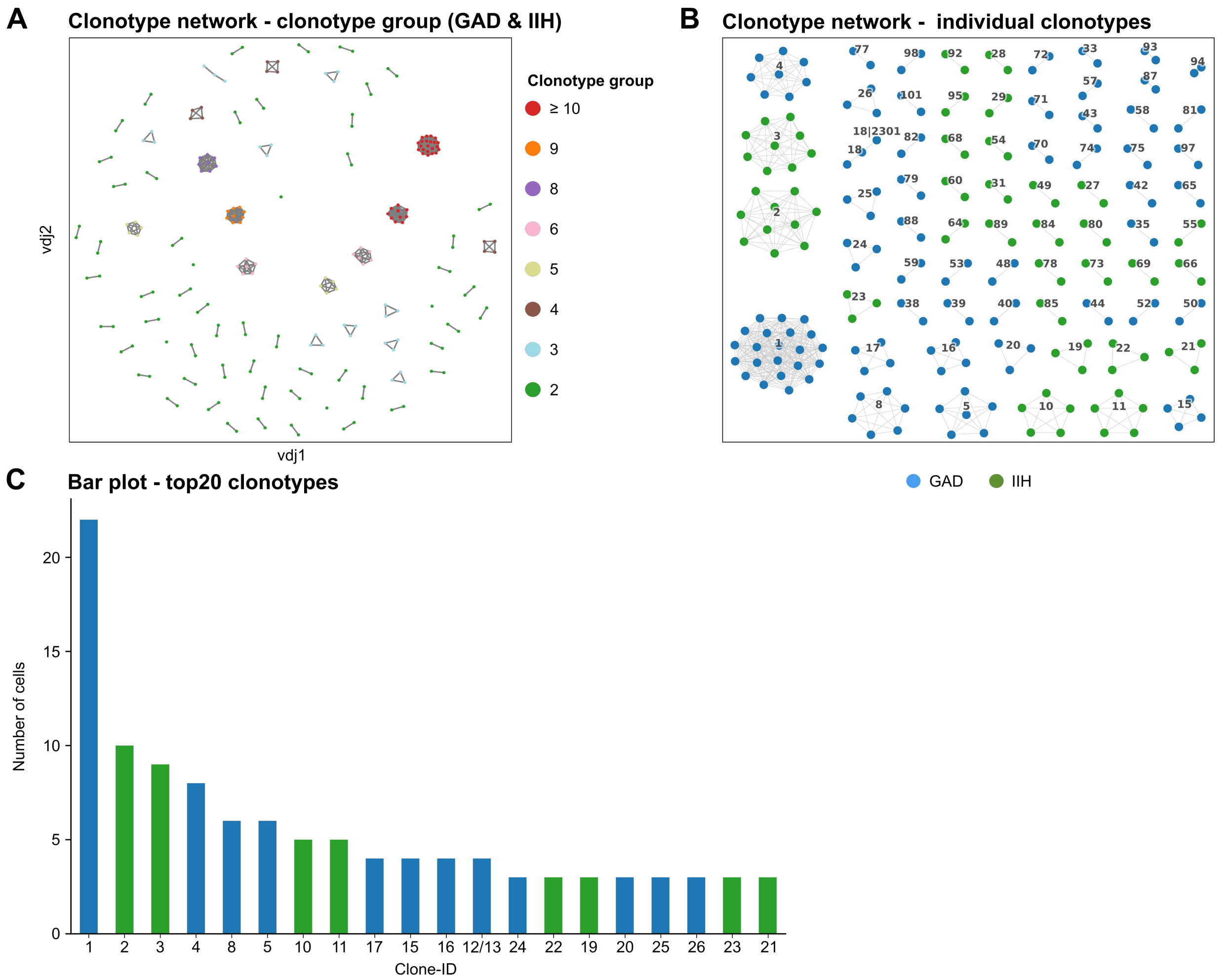


**Supplementary Figure 3 - Clonotype network of CD4$^{+}$ stem cell-like memory T cell repertoire in CSF compartment of anti-GAD65 AINS and IIH patients.**

**A** Detailed clonal network analysis focusing on expanded CD4$^{+}$ TSCM cell clonal groups (frequency ≥ 2 to ≥10) in CSF of anti-GAD65 AINS and IIH patients. **B** Individual clonotype network visualization in CSF, comparing repertoire architecture between anti-GAD65 AINS and IIH patients. Node size reflects clonal frequency. **C** Bar plot represents comparative analysis of the top 20 most abundant CD4$^{+}$ TSCM cell TCR clones in CSF, showing relative frequencies between anti-GAD65 AINS and IIH patients.

*TSCM - stem cell-like memory T cells; CSF - cerebrospinal fluid; GAD - patients with glutamic acid decarboxylase antibody-associated autoimmune neurological syndromes (AINS); IIH - idiopathic intracranial hypertension; TCR - T cell receptor.*
