## Supplementary Table 1 for "CSF single-cell RNA sequencing reveals clonally expanded CD4^+^ stem cell-like memory T cells in GAD65-antibody associated neurological syndromes"

| ID | Analysis | Sample type | Diagnosis | Dominant clinical phenotype | Age | Sex | Disease duration at sampling | Comorbidities | Previous IMT | IMT at time of sampling |
|---|---|---|---|---|---|---|---|---|---|---|
| K_KS_GAD | GEX & BCR & TCR (5' reagent kit) | PBMCs | GAD-CA | CA | 39 | f | 7 Y | DM type 1, s/p breast cancer | IVIG 6 Y pts | No |
| MS-A15 | GEX (3' reagent kit) | CSF | GAD-LE | LE | 59 | f | 4 Y | Hx of thymoma, meningeoma, aHT | Steroids > 6 M pts | No |
| MS-A24 | GEX & BCR & TCR (5' reagent kit) | CSF | GAD-SPS | SPS | 40 | f | 3 Y | Crohn's disease, hypothyroidism (suspected Hashimoto's), aHT | No | No |
| RO-03 | GEX & BCR & TCR (5' reagent kit) | PBMCs & CSF | GAD-LE | LE | 47 | f | 3.5 Y | De novo insulin dependent DM | No | No |
| RO-14 | GEX & BCR & TCR (5' reagent kit) | PBMCs & CSF | GAD-LE | LE | 37 | f | 10.5 Y | None | No | No |
| MS-A18 | GEX (3' reagent kit) | CSF | GAD-LE | LE | 46 | m | Few days | Adrenal-gland incidentaloma, hypokaliemia, AV block I, aHT, obesity, OSAS, degenerative joint pain | No | No |
| MUC-9910 | GEX & BCR & TCR (5' reagent kit) | PBMCs & CSF | GAD-CA | CA | 72 | f | 6 M | Glaucoma, aHT, hypothyreodism | No | No |
| MUC-9961 | GEX & BCR & TCR (5' reagent kit) | PBMCs & CSF | GAD-SPS & -LE | SPS | 58 | f | 3 W | DM type 1, Hashimoto's, Vitiligo, s/p breast cancer | No | No |

**Supplementary Table 1: Basic demographic and clinical data of anti-GAD-AINS patients**

aHT - arterial hypertension; anti-GAD65 AINS patients - patients with glutamic acid decarboxylase antibody-associated autoimmune neurological syndromes; AV - atrioventricular; BCR - B cell receptor; CA - cerebellar ataxia; CSF - cerebrospinal fluid; DM - diabetes mellitus; f - female; GAD - Glutamic acid decarboxylase; GAD-AINS - patients with glutamic acid decarboxylase antibody-associated autoimmune neurological syndromes; GEX - gene expression; Hx - history of; IA - immunoadsorption; ID - identifier; IMT - immunomodulatory therapy; LE - limbic encephalitis; m - male; M - months; OSAS obstructive sleep apnea syndrome; PBMCs - peripheral blood mononuclear cells; PE - plasmapheresis; pts - priot to sampling; Seq - sequencing; s/p - status post; SPS - stiff person syndrome; TCR - T cell receptor; W - weeks; Y - years.
