## Supplementary Table 2 for "CSF single-cell RNA sequencing reveals clonally expanded CD4^+^ stem cell-like memory T cells in GAD65-antibody associated neurological syndromes"

| ID | CSF cells | BCSFBD | Protein (mg/dl) | Serum Alb (g/l) | CSF Alb | Q Alb (x1/1000) | Serum IgG (g/l) | CSF IgG (mg/l) | Q IgG (x1/1000) | Q IgA (x1/1000) | Q IgM (x1/1000) | Ocbs | Serum GAD65-AABs (assay) | CSF GAD65-AABs (assay) | ASI |
|---|---|---|---|---|---|---|---|---|---|---|---|---|---|---|---|
| K_KS_GAD | 1 | No | 417 | n/a | n/a | 6.3 | 15.1 | 45.9 | 3.0 | 2.3 | 2.0 | No | >2000 IU/ml (ELISA) | >2000 IU/ml (ELISA) | n/a |
| MS-A15 | 1 | No | 411 | 43.5 | 162 | 3.1 | 8.1 | 17.1 | 2.1 | 0.7 | 0.2 | No | 1:100 (CBA) | 1:100 (CBA) | 473.0 |
| MS-A24 | 11 | Yes | 603 | 40.4 | 308 | 7.1 | 12.5 | 3.9 | 0.3 | 1.8 | 0.5 | No | 1:100000 (CBA)<br>997000 IU/ml (ELISA) | 1:1000 (CBA)<br>7160 IU/ml (ELISA) | n/a |
| RO-03 | 1 | No | 230 | 45 | 155 | 3.1 | 18.2 | 31 | 1.7 | n/a | n/a | n/a | pos. (IHC)<br>6150 IU/ml (ELISA) | pos. (IHC)<br>12 IU/ml (ELISA) | 4.2 |
| RO-14 | 3 | No | 290 | 44 | 195 | 4.1 | 9.2 | 20 | 2.2 | n/a | n/a | No | pos. (IHC) | neg. (IHC) | 0.9 |
| MS-A18 | 6 | No | 370 | 39.8 | 165 | 4.1 | 8.9 | 54.8 | 6.2 | 1.4 | 0.3 | Yes | 1:1000 (CBA) | 1:100 (CBA) | 35.9 |
| MUC-9910 | 7 | No | 49 | 46.6 | 449 | 9.1 | 1.2 | 16.0 | 13.3 | 4.1 | 1.3 | Yes | 1:800 (CBA) | 1:100 (CBA) | 16.8 |
| MUC-9961 | 3 | No | 32 | 43.3 | 199 | 4.1 | 10.5 | 21.0 | 2.0 | 0.8 | 0.1 | Yes | 1:6400 (CBA) | 1:200 (CBA) | 15.6 |

**Supplementary Table 2: Basic CSF characteristics of anti-GAD65 AINS patients**

AABs - autoantibodies; Alb - albumin; anti-GAD65 AINS patients - patients with glutamic acid decarboxylase antibody-associated autoimmune neurological syndromes; ASI - antibody specificity index; BCSFBD - blood-CSF barrier dysfunction; CBA - cell-based assay; CSF - cerebrospinal fluid; ELISA - Enzyme-linked Immunosorbent Assay; GAD65 - 65 kDa isoform of the glutamic acid decarboxylase; ID - identifier; IHC - immunohistochemistry; Ig - immunoglobulin; n/a - not available; neg . - negative; Ocbs - oligloclonal bands; pos. - positive; Q - serum/CSF ratio
