## Supplementary Table 3 for "CSF single-cell RNA sequencing reveals clonally expanded CD4^+^ stem cell-like memory T cells in GAD65-antibody associated neurological syndromes"

| ID | Analysis | Sample type | Diagnosis | Age | Sex | Comorbidities | Previous IMT | IMT at time of sampling |
|---|---|---|---|---|---|---|---|---|
| MS-MPST83775 | GEX (3' reagent kit) | PBMC & CSF | IIH | 42 | m | None | No | No |
| MS-MPST95809 | GEX (3' reagent kit) | PBMC + CSF | IIH | 43 | f | None | No | No |
| RO-21 | GEX & TCR & BCR (5' reagent kit) | PBMC + CSF | IIH | 35 | f | None | No | No |
| RO-34 | GEX & TCR & BCR (5' reagent kit) | PBMC + CSF | IIH | 36 | f | None | No | No |
| MUC-IIH-8360-KE | GEX & TCR & BCR (5' reagent kit) | PBMC + CSF | IIH | 27 | f | Migraine, mild depressive syndrome | No | No |
| MUC-IIH-8361-KE | GEX & TCR & BCR (5' reagent kit) | PBMC + CSF | IIH | 52 | f | Hypothyroidism, aHT | No | No |
| RO-31-6894844 | GEX & TCR & BCR (5' reagent kit) | PBMC + CSF | IIH | 40 | f | Fibromyalgia | No | No |
| RO-32-4868164 | GEX & TCR & BCR (5' reagent kit) | PBMC + CSF | IIH | 36 | f | None | No | No |

**Supplementary Table 3: Basic demographic and clinical data of IIH patients**

aHT - arterial hypertension; BCR - B cell receptor; CSF - cerebrospinal fluid; f - female; GEX - gene expression; ID - identifier; IIH - idiopathic intracranial hypertension; IMT - immunomodulatory thetrapy; m - male; PBMCs - peripheral blood mononuclear cells;  Seq - sequencing; TCR - T cell receptor.
