## Supplementary Table 4 for "CSF single-cell RNA sequencing reveals clonally expanded CD4^+^ stem cell-like memory T cells in GAD65-antibody associated neurological syndromes"

| ID | CSF cells (/µl) | BCSFBD | Protein (mg/dl) | Serum Alb (g/l) | CSF Alb (mg/l) | Q Alb (x1/1000) | Serum IgG (g/l) | CSF IgG (mg/l) | Q IgG (x1/1000) | Q IgA (x1/1000) | Q IgM (x1/1000) | Ocbs |
|---|---|---|---|---|---|---|---|---|---|---|---|---|
| MS-MPST83775 | 6 | Yes | 873.0 | 38.7 | 396.0 | 10.2 | 7.6 | 35.5 | 4.6 | 3.3 | 0.5 | No |
| MS-MPST95809 | 1 | No | 277.0 | 39.1 | 132.0 | 3.4 | 13.4 | 20.6 | 1.5 | 1.8 | 0.2 | No |
| RO-21 | 3 | No | 390.0 | 39.0 | 239.0 | 6.1 | 10.1 | 34 | 3.4 | n/a | n/a | No |
| RO-34 | 1 | n/a | 130.0 | n/a | 78.0 | n/a | 11.8 | 13 | 1.1 | n/a | n/a | No |
| MUC-IIH-8360-KE | 2 | n/a | 320.0 | n/a | n/a | n/a | n/a | n/a | n/a | n/a | n/a | n/a |
| MUC-IIH-8361-KE | 2 | n/a | 490.0 | n/a | n/a | n/a | n/a | n/a | n/a | n/a | n/a | n/a |
| RO-31-6894844 | 1 | n/a | 370.0 | 39.0 | n/a | n/a | n/a | n/a | n/a | n/a | n/a | n/a |
| RO-32-4868164 | 10 | n/a | 210.0 | n/a | n/a | n/a | n/a | n/a | n/a | n/a | n/a | n/a |

**Supplementary Table 4: Basic CSF characteristics of IIH patients**

Alb - albumin; BCSFBD - blood-CSF barrier dysfunction; CSF - cerebrospinal fluid; ID - identifier; Ig - immunoglobulin; IIH - idiopathic intracranial hypertension; n/ - not available; Ocbs - oligloclonal bands; Q - serum/CSF ratio.
